# Mapping the ISR Landscape in Cognitive Disorders *via* single-cell multi-omics

**DOI:** 10.1101/2025.02.28.640905

**Authors:** Kristof Torkenczy, Lucas C. Reineke, Sean W. Dooling, Benjamin Henderson, Daniel N. Itzhak, Benjamin Yang, Dongze He, Richard M. Myers, Peter Walter, Stefka Tyanova, Mauro Costa-Mattioli

**Affiliations:** Altos Labs, Inc., Bay Area Institute, Redwood City, California, 94665, USA; HudsonAlpha Institute for Biotechnology, Huntsville, Alabama, 35806, USA; Department of Neuroscience, Baylor College of Medicine, Houston, Texas, 77030, USA

**Keywords:** proteostasis networks, single-cell RNA-sequencing, single-cell ATAC-sequencing, proteomic, cognitive decline, integrated stress response

## Abstract

Persistent activation of the integrated stress response (ISR) is a major driver of cognitive decline in both neurodevelopmental and neurodegenerative disorders. Using a new mouse model (*Ppp1r15b*^R658C^ mice) that mimics the persistent ISR activation and cognitive decline observed in humans, we generated the first single-cell ISR atlas of the brain. By integrating single-cell RNA-seq and single-cell ATAC-seq with proteomics, we discovered that distinct brain cell types respond differently to persistent ISR activation and elicit cell-type-specific ISR programs. Interestingly, chromatin accessibility analyses revealed that the ISR downstream factor ATF4 is a key ISR effector in GABAergic neurons, while AP-1 (JUNB) is implicated in glutamatergic neurons. More importantly, selective deletion of ATF4 in GABAergic neurons—but not in glutamatergic neurons—impacts ISR-mediated cognitive decline in *Ppp1r15b*^R658C^ mice, demonstrating that different neuronal subtypes rely on unique ISR downstream effectors to regulate mnemonic processes. Furthermore, we defined a comprehensive molecular signature of persistent ISR activation, which we showed could serve as a biomarker for cognitive dysfunction across neurodevelopmental, neurodegenerative disorders and normal aging. This multi-omic framework provides a key platform for exploring and validating new scientific hypotheses, significantly advancing our understanding of ISR-related brain disorders.

## Introduction

The integrated stress response (ISR) is a conserved signaling network that restores protein homeostasis by regulating protein synthesis. In the brain, the ISR acts as a “memory rheostat”, fine-tunning cognitive processes in health and disease^1^. ISR activity modulates both long-lasting increases and decreases in synaptic function^2–4^, two key forms of synaptic plasticity thought to be important for learning and memory^5^. Extensive genetic and pharmacological evidence demonstrates that while activation of the ISR impairs long-term memory, ISR inhibition facilitates memory consolidation^2,3,6–14^. Furthermore, while acute ISR activation can be protective in neurons, persistent ISR activation has been causally linked to the cognitive dysfunction observed in a wide range of disorders ranging from rare genetic conditions^15–20^ to common neurodevelopmental and neurodegenerative diseases, such as Down syndrome^21^ and Alzheimer’s disease^22–24^, respectively, as well as age-related cognitive decline^10,25^. Consequently, ISR inhibition is emerging as a promising strategy for reversing cognitive dysfunction in a wide range of memory disorders resulting from disruption in protein homeostasis^1^.

Despite its pivotal role in cognitive function, the precise mechanisms by which persistent ISR activation leads to long-term memory deficits remain unclear. Cognitive function relies on the coordinated activity of specialized neuronal and glial cells. Therefore, dissecting the ISR’s role at the cellular level is essential. Identifying which cell populations are most vulnerable to persistent ISR activation—and how these vulnerabilities contribute to memory impairment—will help uncover the cellular and molecular basis of cognitive decline. Several key questions arise: Do neurons and glial cells respond to ISR activation in similar ways, or are there distinct patterns of activation in different cell types? Do different brain cells utilize the same downstream effectors, or do they employ unique mechanisms? Can a common ISR signature be identified that distinguishes healthy from pathological states?

To address these questions, and given that the ISR inhibits general translation while promoting the synthesis of specific transcription factors that reprogram gene expression, such as ATF4^1,26,27^, we employed a comprehensive multi-omic strategy. This approach, which integrates single-cell RNA sequencing (scRNA-seq) and single-cell assay for transposase-accessible chromatin sequencing (sc-ATAC-seq) with bulk proteomics and phosphoproteomics, not only provides new insights into the molecular and cellular mechanisms underlying ISR-driven cognitive impairments, but also paves the way for targeted therapies to treat memory-related disorders where the ISR plays a central role.

Specifically, we developed a novel model of persistent ISR activation driven by a human genetic variant that closely mimics the cognitive decline observed in patients (see accompanying manuscript by Reineke *et al.*). This model allows precise interrogation of ISR-driven gene expression changes in specific cell types and brain regions. Our multi-omic integration platform unexpectedly revealed that the ISR impacts different cell types in distinct ways, challenging the notion of a uniform ISR response across brain cells. Further molecular genetics studies revealed that different cell types rely on unique ISR effectors to regulate long-term memory. Finally, we defined a molecular signature of persistent ISR activation, which serves as a biomarker of cognitive impairment across neurodevelopmental and neurodegenerative disorders, including aging. This multi-omic approach lays a powerful foundation for formulating and evaluating new hypotheses, thereby enhancing our understanding of neurological disorders associated with ISR activation.

## Results

### Distinct patterns of ISR activation across different brain cell types

To understand how neural circuits give rise to different behavioral states, it is important to explore the cell types that compose these circuits and their specific roles in processing and integrating information. Recent advances in high-throughput single-cell sequencing technology have allowed the identification of several neuronal and glial cell types based on their transcriptional profile^28,29^. However, the impacts of the ISR in each of these cell type populations remains poorly understood. To answer this question and construct an ISR brain atlas, we employed a new mouse model that carries a human mutation in a central component of the ISR (*Ppp1r15b*^R658C^ mice; see accompanying paper by Reineke *et al*.). These mice exhibit a selective and persistent activation of the ISR in the brain, resulting in impaired long-lasting synaptic plasticity and long-term memory, closely mirroring the human disease phenotype. Importantly, inhibiting the ISR in this model reverses the deficits in synaptic plasticity and long-term memory, indicating that these impairments are directly attributed to ISR activation. Thus, we performed single-cell (10X) multi-omic sequencing across cell types from the brains of WT and *Ppp1r15b*^R658C^ mice (**Figure 1A**). Briefly, nuclei were extracted from cortical tissue to simultaneously perform scRNA-seq and scATAC-seq and assess both gene expression changes and chromatin accessibility across brain cells, respectively (**Figure 1B**). After filtering of the data, we retained 85,402 high-quality nuclei covering 28,601 genes and 308,774 *cis*-regulatory elements (CREs; **Figure S1**). Notably, the data set exhibited a high number of unique reads, with minimal contamination from mitochondrial genes, and a high number of detected genes per cell (see **Figure S1**). In addition, we observed a significant number of shared accessible peaks, strong enrichments at transcription start sites (TSS), and high nucleosome signal strength across cells and samples (**Figure S1A-S1B**), demonstrating the high quality of the scATAC-seq data, which provides a robust foundation for in-depth analyses.

**Figure 1.**
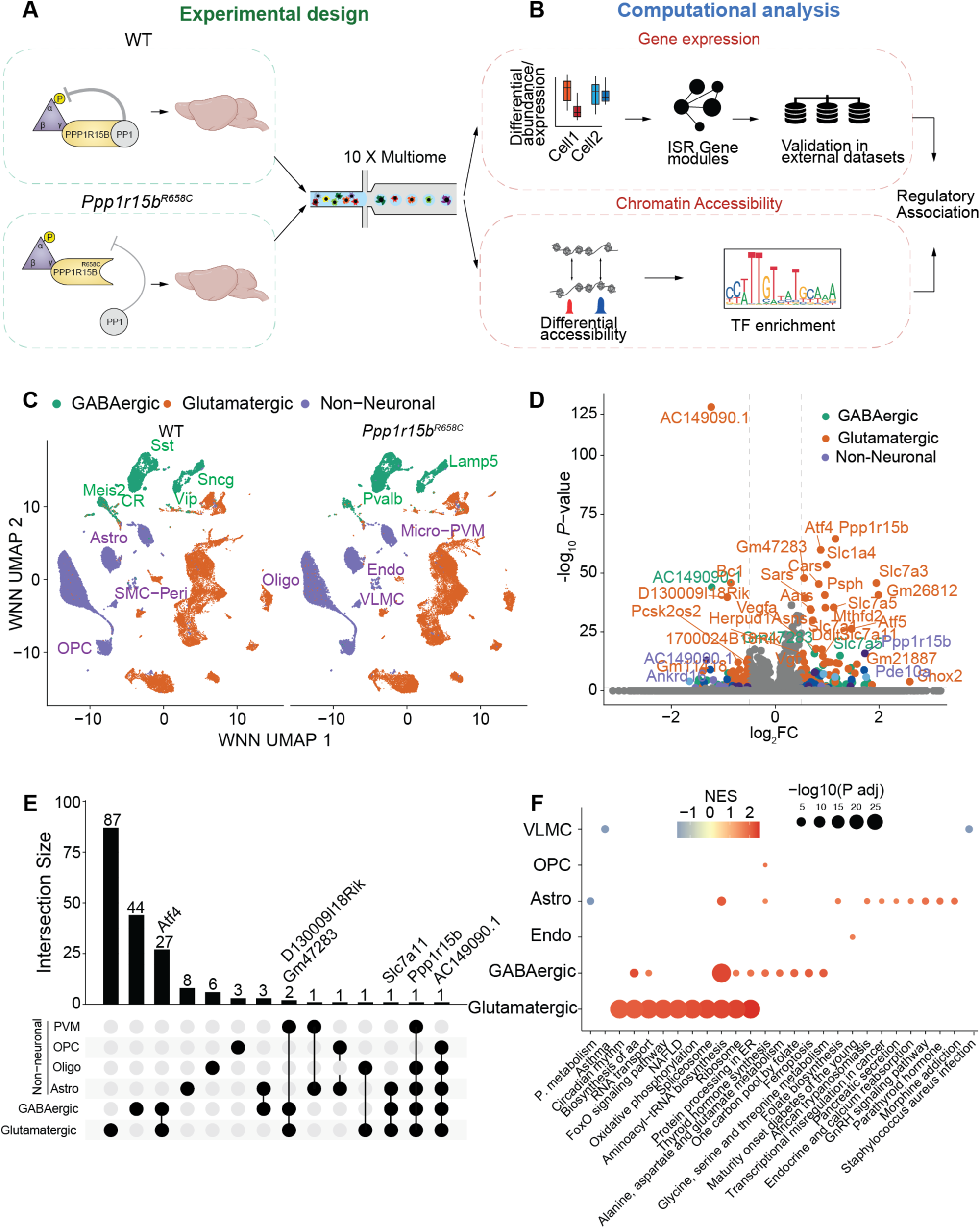
Single cell multi-omic map of persistent ISR activation. (A) Overview of the experimental design to map the single-cell multi-omic landscape of brain cell populations in mice with persistent ISR activation (*Ppp1r15^R658C^*) compared to control (WT) mice (*n* = 7 per group). (B) Schematic of the computational analysis pipeline used to identify single-cell differences in gene expression and chromatin accessibility between brain samples from WT and *Ppp1r15^R658C^* mice. (C) Joint weighted UMAP projections of 85,502 cells from WT (left) and *Ppp1r15^R658C^*(right) mice, visualized across the three major cell classes: GABAergic, glutamatergic and non-neuronal lineages. (D) Volcano plot of differential gene expression analysis between WT and *Ppp1r15^R658C^* brain cells. Differentially expressed genes (marked by dashed lines) with adjusted *P*-value < 0.05 and log2_-_ fold change > 0.25 are categorized by GABAergic, glutamatergic and non-neuronal lineages. (E) Cell type-specific unique and overlapping differentially expressed genes between brain cells from WT and *Ppp1r15b*^R658C^ mice. (F) KEGG pathway gene set enrichment analysis of differentially expressed genes in each cell type of brain samples from WT and *Ppp1r15*^R658C^ mice. Significant pathways were selected based on adjusted *P*-value threshold of < 0.05. Normalized enrichment score (NES) and -log10 adjusted *P*-value are represented by the color and size of the dots, respectively.

We next performed principal component analysis (PCA) for gene expression, and latent semantic analysis (LSI) for chromatin accessibility^30,31^. By reference mapping to the Allen Brain Atlas^32^ and manual annotation based on known cell-types markers, we identified broad brain cell types. We found that the overall cell type composition was not changed, as determined by the weighted-nearest neighbor (WNN) Projection of data using Uniform Manifold Projection (UMAP; **Figure 1C** and **Figure S1C-S1E**), in the brains of *Ppp1r15b*^R658C^ mice. Strikingly, ISR-induced genes exhibited varying degrees of up-regulation depending on the cell type in *Ppp1r15b*^R658C^ mice (**Figure 1D**). Moreover, we found no genes with ubiquitous regulation across all brain cell type despite persistent ISR activation (**Figure 1E**). It is noteworthy that despite the increase in its mRNA level, PPP1R15B protein levels were not changed in the brain of *Ppp1r15b*^R658C^ mice (see accompanying manuscript by Reineke *et al*.). In contrast, most genes in mice with persistent ISR activation were expressed in a cell type specific manner: for example, 27 genes (including *Atf4*) were shared between glutamatergic and GABAergic neurons, 43 were specific to glutamatergic neurons, and 17 were specific to GABAergic neurons. Hence, each cell type responds to the ISR by modulating different subsets of genes (**Figure 1E-F**).

To investigate how persistent ISR activation impacts different cell types, we first quantified the levels of *Atf4* —a key transcription factor and major ISR downstream target^1^—in each cell type as a proxy for ISR activity. Since ISR activation triggers ATF4 translation^26,27^, its expression level can reflect ISR activation. In *Ppp1r15b*^R658C^ mice, we found that *Atf4* expression was significantly higher in various cell types, including glutamatergic and GABAergic neurons, whereas oligodendrocytes showed increased *Atf4* expression but no changes in its downstream targets (e.g., *Atf5, Atf6* and *Slc7a11,* **Figure S2A**). To further investigate the cell type-specific effects of persistent ISR activation, we performed gene set enrichment analysis (GSEA) for each cell type using the KEGG database, ranking genes by the magnitude and significance of their differential expression. We found that persistent ISR activation led to distinct gene set enrichment patterns in each cell type, highlighting the cell type-specific nature of ISR activation (**Figure 1F**). When we expanded our analysis to include molecular function and biological processes, we also observed similar cell type-specific changes upon persistent ISR activation (**Figure S2B-C**). These findings highlight that persistent ISR-driven gene expression is highly cell type-dependent, suggesting that distinct ISR downstream effectors mediate unique functional outcomes across different cell types.

### Identification of an ISR signature using scRNA-seq: a potential biomarker for cognitive disorders

While ATF4 target gene expression is often used as a surrogate marker for ISR activation, it is important to note that ATF4 expression can also be modulated by ISR-independent mechanisms. For instance, activation of the mechanistic target of rapamycin (mTORC1) promotes *Atf4* mRNA translation^33–35^ in an ISR-independent manner, and ATF4 levels are modulated by methylation of the ATF4 promoter, post-translational modifications and protein degradation^36^. Consequently, defining a robust ISR signature at the single-cell level requires a broader approach that extends beyond just ATF4 and its downstream targets. To achieve this and identify a comprehensive ISR signature, we employed metacell-based approaches, which cluster similar cells based on gene expression patterns, thereby improving the signal-to-noise ratio in single-cell genomics^37^ (**Figure S3A-B**). Initially, we used MiloDE^38^, a recently developed statistical framework designed to detect subtle changes in gene expression that are often undetectable with other traditional single-cell methods. Using this approach, we identified neighborhoods of cells with significant changes in genes expression across the brain of WT and *Ppp1r15b*^R658C^ mice (**Figure 2A**).

**Figure 2.**
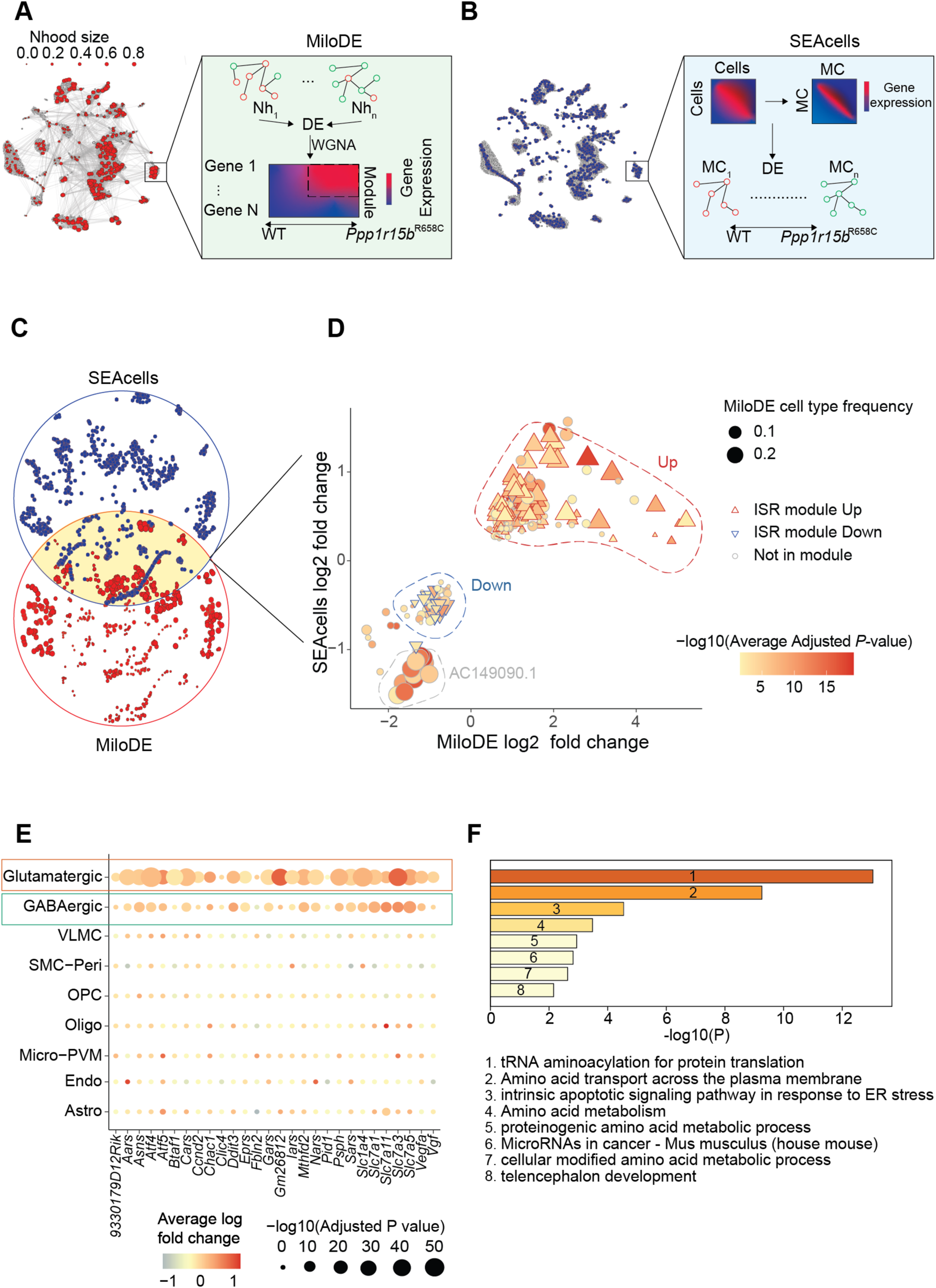
Defining an ISR-specific gene signature. (A) Computational workflow for identifying ISR-related gene modules using MiloDE. Left panel shows the identified cell neighborhoods with neighborhood size, and right panel provides an overview of the analysis approach (B) Computational pipeline for identifying ISR related gene modules using SEAcells. Left panel displays identified metacells, and right panel provides an overview of the analysis approach. (C) Framework for defining the persistent ISR gene signature. (D) Comparison of gene modules derived from MiloDE and SEAcells analyses. Differentially expressed genes (DEGs) between *Ppp1r15b*^R658C^ and WT mice are categorized into ISR up- and down-modules. Log2-fold change values for both SEAcells and MiloDE analyses are shown, with adjusted *P*-values indicated. The relative frequency of DEGs is represented by the size of the shapes, with gene membership to the up- and down-modules indicated by shape type. (E) Log2-fold and adjusted *P*-value of up regulated ISR gene co-expression module identified by SEAcells. (F) Metascape analysis of the ISR *up*-module, showing the top 8 enriched ontologies.

Next, we applied single-cell weighted gene co-expression network analysis (scWGCNA) to these cell neighborhoods to identify gene modules that are up- and down-regulated in the brain of *Ppp1r15b*^R658C^ mice compared to control mice. To ensure the robustness of the identified modules, we iteratively tested various thresholds for differential expression (DE) within cell neighborhoods (**Figure S4**). We identified two highly reproducible gene modules, each containing approximately 30 genes (**Figure S4C-D**). One of these modules, which we termed the *ISR up-*module, showed increased expression of a shared set of ISR genes in several cell types, including glutamatergic and GABAergic neurons, oligodendrocytes, and astrocytes. However, the magnitude of the response varied between cell types (**Figure S4C, S5A-B**). The second module, the *ISR-down* module, included primarily ISR-modulated genes whose expression is reduced in glutamatergic and GABAergic neurons (**Figure S4D, S5C-D**). Both the *ISR up-* and *down-*modules were more enriched in cortical layers, where excitatory and inhibitory neurons are abundant. In contrast, other cell types, such as endothelial cells, did not show consistent differential expression of these genes (**Figure S5A-D**). Moreover, we also identified a gene module that was up-regulated only in oligodendrocytes (**Figure S5E-F**), further supporting the notion that distinct brain cell types respond differently to persistent ISR activation.

To refine the ISR signature, we next employed SEACells^39^, a method that groups single cells with similar expression profiles into high-quality metacells (**Figure 2B**). This method detects subtle changes in low expressed genes while preserving biological variability across cell types. After building metacells with SEACell, we projected them into the same UMAP space as MiloDE, identifying a similar distribution across cell types. We then compared the differentially expressed genes identified by SEACells and MiloDE and established a consensus ISR signature (**Figure 2C-D**). Briefly, we detected 28 genes from the *ISR up-*module (**Figure 2E**). Although we also detected a gene set that was downregulated (ISR *down*-module) under persistent ISR activation (**Figure 2D**), many of the genes in this set were mouse specific, limiting its utility for comparison with human datasets. As a result, we defined the ISR *up*-module as the definitive *“ISR signature”* for further validation.

To further characterize the ISR signature, we performed gene ontology analysis. We found significant changes in pathways involved in tRNA aminoacylation (e.g. *Aars*, *Eprs*, *Nars*, *Sars)*, amino acid transport (e.g. *Slc1a4*, *Slc7a11*, *Slc7a3*, *Slc7a5*) and amino acid metabolism (e.g. *Psph*, *Chac1*, *Mthfd2*; **Figure 2F**), indicating that the ISR signature reflects changes in protein synthesis-related processes and metabolism. Notably, about one thirds of the genes in the ISR signature have not been previously identified as direct targets of ATF4. Thus, the ISR signature defined here provides a more comprehensive and precise representation of ISR activation, which is not exclusively driven by ATF4 regulation, which, as mentioned above, can also be induced in an ISR-independent manner (ref).

Given that the ISR is activated in various cognitive disorders associated with both neurodevelopmental and neurodegenerative conditions^1^, we aimed to determine whether this signature could serve as a reliable biomarker for these disorders (**Figure 3A**). First, we analyzed human single-cell data from Down Syndrome^40^, the leading cause of intellectual disability and a neurodevelopmental disorder^41^ where ISR activation is causally linked to the synaptic and cognitive deficits observed in a mouse model of the disease^21^ (see also accompanying manuscript, Reineke *et al.*) Intriguingly, we found upregulation of the ISR genes across all cell types examined from individuals with Down Syndrome (**Figure 3B** and **Figure S6A**). Next, to validate the ISR signature in neurodegenerative disorders, we analyzed single-cell datasets from human individuals with either Lewy Body Dementia or Parkinson’s disease^42^. In both neurodegenerative disorders, the ISR signature was upregulated (**Figure 3C-D**), although the extent and affected cell types varied between the two diseases. Specifically, in Parkinson’s disease, but not in Lewy Body Dementia, the ISR signature was detectable in astrocytes (**Figure S6B-C**), whereas neurons in both diseases showed enrichment of the ISR signature.

**Figure 3.**
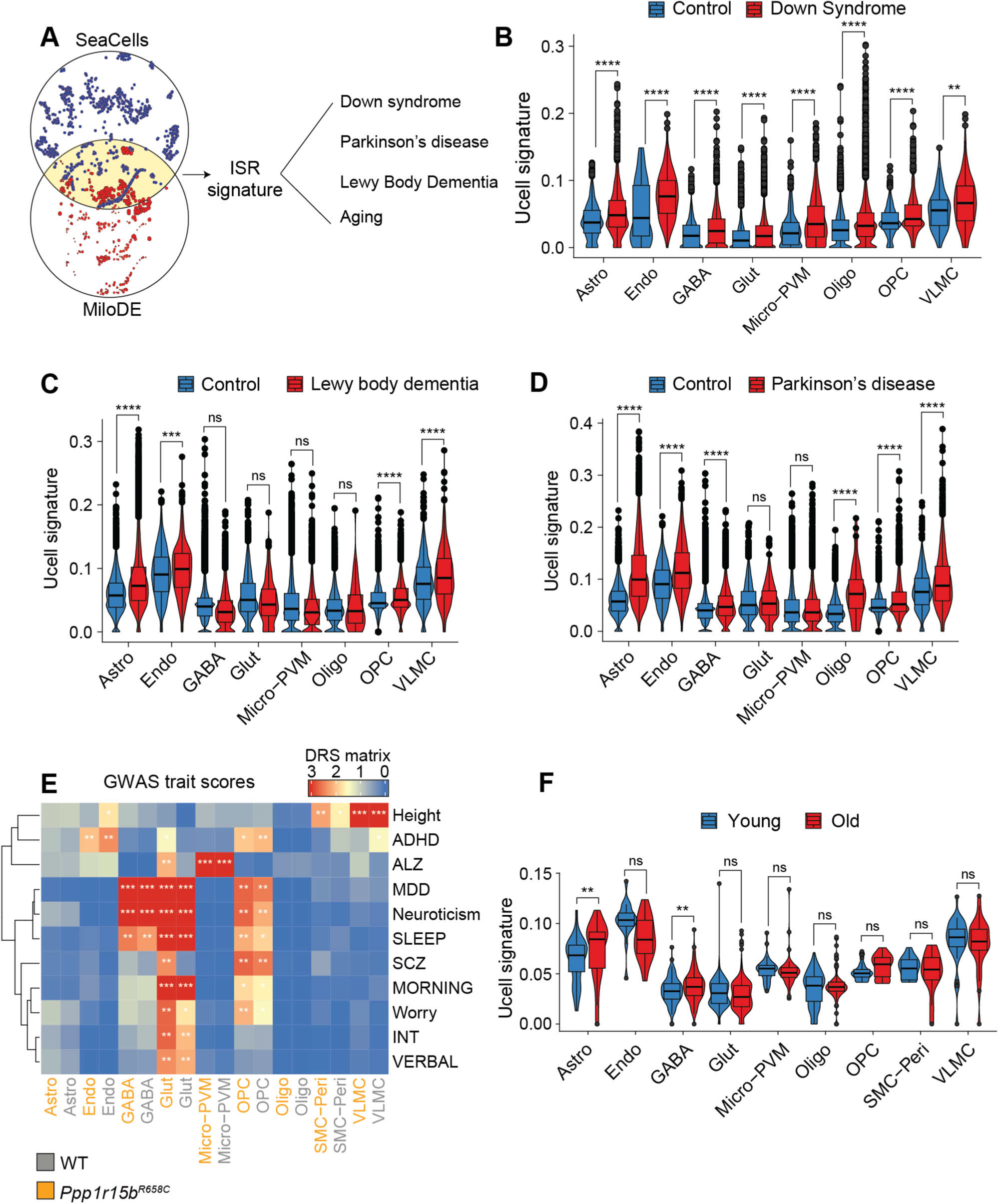
The ISR gene signature is present in diseases associated with persistent ISR. (A) Schematic showing the selection of genes included in the ISR gene signature, and application to diseased datasets. (B) Ucell scores of the ISR gene signature in control and Down Syndrome individuals across cell types (\*\**P* < 0.01, \*\*\*\**P* < 0.0001). (C) Ucell scores of the ISR gene signature in control and individuals with Lewy body dementia across cell types (\*\*\**P* < 0.001, \*\*\*\**P* < 0.0001). (D) Ucell scores of the ISR gene signature in control and Parkinson’s Disease individuals across cell types (\*\*\*\**P* < 0.0001). (E) Comparison of the ISR gene signature with genes identified in human genome-wide association studies in the indicated diseases. ADHD, attention deficit/hyperactivity disorder; ALZ, Alzheimer’s; MDD, major depressive disorder; Neuroticism; Sleep, sleep deprivation; SCZ, schizophrenia; Morning; Worry; INT, intelligence; Verbal, verbal numeric reasoning. (F) Ucell scores of the ISR gene signature in young and old mice across cell types (\**P* < 0.05).

Moreover, using single-cell disease relevance scoring^43^, a method that compares scRNA-seq data with genome-wide association studies, we found the ISR signature was not associated with traits including height and anxiety but successfully revealed differences in glutamatergic neurons in both Alzheimer’s disease and Schizophrenia (**Figure 3E**). While ISR activation is known to contribute to the cognitive decline in Alzheimer’s disease^22–24^ (see also accompanying manuscript, Reineke *et al*.), its possible involvement in schizophrenia is, to our knowledge, a novel finding.

Finally, given that several of these human diseases are age-related, we investigated whether the ISR signature could be detected during aging. Remarkably, we found that the ISR up-signature was present in astrocytes and GABAergic neurons in aged mice (**Figure 3F**). Collectively, these findings suggest that the ISR signature could serve as a biomarker for cognitive decline in both neurodevelopmental and neurodegenerative disorders, with potential implications for understanding the underlying mechanisms of these diseases.

### Cell type-specific roles of ISR downstream effectors during persistent ISR activation

To better understand how the ISR regulates gene expression at the single-cell level, we next examined the association between chromatin accessibility and transcriptional changes during persistent ISR activation. Using scATAC-seq in WT and *Ppp1r15b*^R658C^ mice, we explored global chromatin dynamics (**Figure 1B**). We first investigated the behavior of DNA-binding proteins *via* the chromatin accessibility tool chromVAR^44^. This method calculates the genome-wide chromatin accessibility at loci containing DNA-binding motifs and computes a Z-score for each individual cell, which is then aggregated by cell type. The Z-score quantifies the deviation from baseline, providing an estimate of putative transcription factor (TF) binding to its motifs. This analysis revealed cell type specific differences in TF motifs enrichment across cell types under persistent ISR activation. Specifically, in *Ppp1r15b*^R658C^ mice, activator protein1 (AP-1) motifs, including FOS and JUN, were enriched in glutamatergic neurons. AP-1, a dimeric complex of basic leucine zipper (bZIP) proteins from the JUN and FOS families, is known for its role in regulating gene expression in response to various stimuli ^45^. Conversely, ATF4 motifs were enriched in GABAergic neurons, and neither ATF4 nor AP-1 binding motifs were significantly enriched in non-neuronal cell subtypes under persistent ISR activation (**Figure 4A**). These findings underscore the distinct mechanisms in which different cell types reprogram gene expression upon persistent ISR activation, further underscoring the cell type-specific roles of ISR downstream effectors.

**Figure 4.**
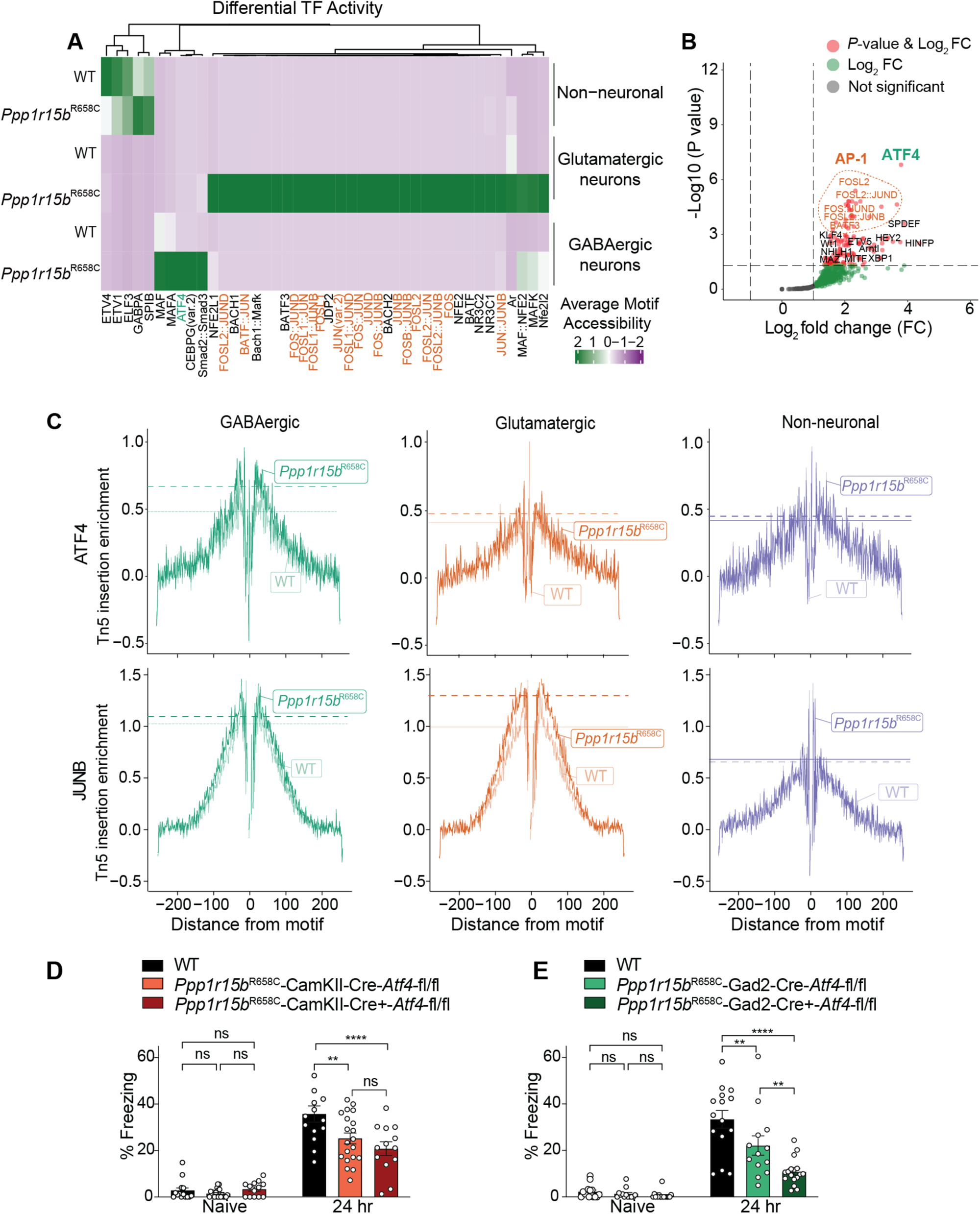
Epigenetic characterization of cell type-specific ISR regulation. (A) Transcription factor activity across cell types and conditions; ChromVar deviation Z-scores are shown. ATF4 and AP-1 members are highlighted in green and orange, respectively. (B) Volcano plot of top transcription factor motif enrichment in ATAC-seq peaks associated with gene expression changes (*P*-value < 0.001 & z-score > 0). Hypergeometric test was performed using a set of GC-content matched background peaks. (C) Bias corrected ATAC-seq footprinting signal centered around ATF4 (top row) and JUNB (bottom row) motifs, aggregated from peaks linked to gene expression changes across cell types. Dotted horizontal lines show *Ppp1r15b*^R658C^ and WT flank height for comparison^30,31^. (D) Contextual long-term fear memory in WT (*n* = 14), *Ppp1r15b*^R658C^*-CamKII-Cre^-^-Atf4^flox/flox^* (*n* = 20), and *Ppp1r15b*^R658C^*-CamKII-Cre^+^-Atf4 ^flox/flox^* (*n* = 13) mice (*F*_2,88_ = 8.78; WT vs. *Ppp1r15b*^R658C^-*CamKII-Cre^-^-Atf4^flox/flox^, t* = 4.47, *P* < 0.01; *Ppp1r15b*^R658C^*-CamKII-Cre^-^-Atf4^flox/flox^*vs. *Ppp1r15b*^R658C-^*CamKII-Cre^+^-Atf4^flox/flox^, t* = 1.55, *P* > 0.99; WT vs. *Ppp1r15b*^R658C^*-CamKII-Cre^+^-Atf4^flox/flox^, t* = 5.48, *P* < 0.0001). (E) Contextual long-term fear memory in WT (*n* = 18), *Ppp1r15b*^R658C^*-Gad2-Cre^-^-Atf4^flox/flox^*(*n* = 13), and *Ppp1r15b*^R658C^*-Gad2-Cre^+^-Atf4^flox/flox^* (*n* = 17) mice (*F*_2,94_ = 8.80; WT vs. *Ppp1r15b*^R658C^*-Gad2-Cre^-^-Atf4^flox/flox^, t* = 3.05, *P* < 0.01; *Ppp1r15b*^R658C^*-Gad2-Cre^-^-Atf4^flox/flox^* vs. *Ppp1r15b*^R658C^*-Gad2-Cre^+^-Atf4^flox/flox^, t* = 3.61, *P* < 0.01; WT vs. *Ppp1r15b*^R658C^*:Gad2-Cre^+^-Atf4^flox/flox^, t* = 7.21, *P* < 0.0001).

To determine whether selective, cell type-specific chromatin opening has functional consequences, we utilized Signac^30,31^, a comprehensive tool for scATAC-seq analyses that links locally accessible chromatin regions to their regulated gene products. Using open chromatin regions associated with differentially expressed genes, we performed TF motif identification. This analysis revealed ATF4-specific enrichment in GABAergic neurons and AP-1 enrichment in glutamatergic neurons (**Figure 4B** and **Figure S7C**). The cell type-specific pattern is further exemplified by two genes, *Gars* and *Fbln2*. *Gars* is differentially expressed and exhibits chromatin opening specifically in GABAergic neurons, while *Fbln2* shows differential expression and chromatin opening in glutamatergic neurons, providing additional evidence for cell type-specific activation of TF motifs in response to persistent ISR activation (**Figure S7A-B**).

To further explore cell type-specific transcriptional regulation, we applied TF footprinting analysis for ATF4 and AP-1 components. This method estimates TF location by measuring the interference of TF binding with Tn5 (transposase) accessibility. When a TF binds to its cognate motif, it prevents Tn5 incorporation at the site, but chromatin surrounding the TF binding site remains accessible, allowing Tn5 to bind. By averaging these measurements across the entire genome and within individual cell types, one can estimate the regulatory relevance of a specific TF in driving gene expression changes. Our results showed enhanced chromatin accessibility for ATF4 in GABAergic neurons and for AP-1 (JUNB) in glutamatergic neurons (**Figure 4C**). We further identified selective enrichment for the DNA-binding motifs of JUNB and FOSL2 in glutamatergic neurons (**Figure S7D**), which are distinct from those of other AP-1 components (**Figure S8A**). Similar results were obtained using a multi-modal tool called Pando^46^, which integrates scRNA-seq and scATAC-seq data to infer gene regulatory networks while adjusting for TF expression levels (**Figure S9**). These results suggest that upon persistent ISR activation specific TFs regulate gene expression in a cell-type specific manner.

The intriguing segregation of ATF4 and AP-1 TF activity to GABAergic and glutamatergic neurons, respectively, under persistent ISR conditions, prompted us to investigate the functional relevance of these findings in cognitive decline caused by persistent ISR activation. Specifically, we assessed whether conditional deletion of ATF4 from glutamatergic or GABAergic neurons differentially affects long-term memory in *Ppp1r15b*^R658C^ mice, a model for persistent ISR activation. To answer this question, we conditionally delete *Atf4* from specific neuronal subtypes. Briefly, *Ppp1r15b*^R658C^ mice were first crossed with *Atf4^flox/flox^* mice, and their progeny further crossed with either a GABAergic-specific (*Gad2-Ires-Cre*)^47^ or a glutamatergic-specific Cre mouse line (*CaMKII-Cre*)^48^ to delete ATF4 in these respective neuronal subtypes, respectively.

We assessed long-term memory using a contextual fear conditional paradigm, where a context (conditioned stimulus) is paired with a foot shock (unconditioned stimulus). Freezing behaviors were measured twenty-four hours after training, as an index of the strength of their long-term memory upon exposure to the context. Consistent with our previous results (see also accompanying manuscript, Reineke *et al.*), *Ppp1r15b*^R658C^ mice with persistent ISR activation showed impaired contextual long-term memory (**Figure 4D**). Interestingly, deleting *Atf4* from glutamatergic neurons from *Ppp1r15b*^R658C^ mice did not significantly affect their long-term memory (*Ppp1r15b*^R658C^*-Atf4^flox/flox^-Cre^CaMKII+^*mice, **Figure 4D**). In contrast, deletion of *Atf4* in GABAergic neurons from *Ppp1r15b*^R658C^ mice (*Ppp1r15b*^R658C^*-Atf4^flox/flox^-Cre^Gad2+^*mice) further exacerbated their long-term memory deficits (**Figure 4E**), indicating that ATF4 has a protective role. Hence, the results highlight that distinct ISR downstream effectors play unique roles across different cell types, demonstrating that ATF4 is only involved in long-term memory in GABAergic neurons during persistent ISR activation.

### A multi-omic fingerprint of persistent ISR activation

Given the ISR regulates gene expression at multiple levels, including translation and transcription^1^, the integration of different omic-modalities could provide a novel perspective on how the ISR regulates mnemonic processes. To investigate ISR-mediated changes at the protein level, we performed proteomic and phosphoproteomic analyses on cortical tissue from WT and *Ppp1r15b*^R658C^ mice. A total of 9,621 proteins were detected (**Figure 5A**). Notably, 21 out of the 28 ISR signature gene products were detected by mass spectrometry, with half of them upregulated (**Figure 5B**). Among the upregulated proteins, we identified members of the tRNA-biosynthetase family (AARS1, CARS1, GARS1, IARS1, NARS1, SARS1; **Figure 5A, C-D**), as well as amino acid transporters (SLC7A1, SLC7A3, SLC7A5, SLC3A2 and SLC7A11), which are known to be induced transcriptionally by ATF4^35,49^. Additionally, MTHFD2, a key protein involved in metabolism and a target of ATF4^33^ also changed at the mRNA level, was significantly upregulated in the brain of *Ppp1r15b*^R658C^ mice (**Figure 5A**, 5D). Consistent with the RNA signature, the proteomic signature extends beyond ATF4 targets, as demonstrated by the increased protein levels of NIBAN1 and CYB5R1 (**Figure 5A**). However, not all changes at the mRNA level are reflected at the protein level. For instance, while EPRS1, the glutamate-tRNA ligase was significantly upregulated at the transcriptome level, EPRS1 protein levels were not different between WT and *Ppp1r15b*^R658C^ mice (**Figure 5C**). These findings underline the necessity of a more comprehensive readout to assess ISR activity than ATF4 transcriptomics.

**Figure 5.**
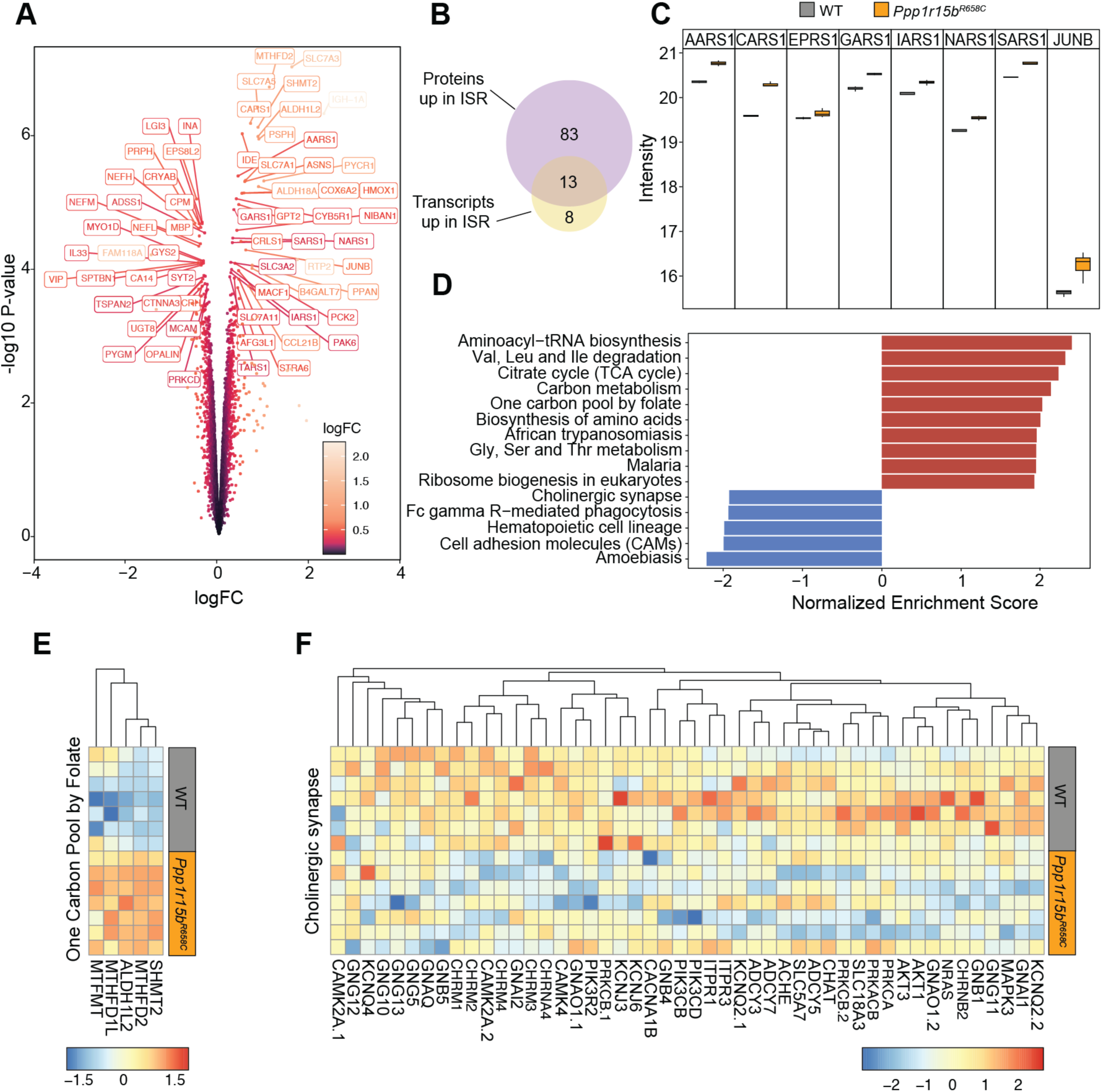
Proteomic characterization of persistent ISR regulation. (A) Volcano plot showing differential protein expression between brain samples from WT and *Ppp1r15b*^R658C^ mice. Proteins are color-coded based on fold change. (B) Comparison of upregulated proteins with the upregulated transcripts from the ISR *up*-signature derived from scRNA-seq of *Ppp1r15^R658C^* mice. Note that, while there are 28 genes in the ISR signature, only 21 were detected in mass spectrometry. (C) Quantification of select proteins, including constituents of the single-cell ISR signature. (D) KEGG gene set enrichment analysis of proteomic data from brain samples from WT and *Ppp1r15b*^R658C^ mice. (E) Heatmap showing components of the “One carbon pool by folate” KEGG gene set. (F) Heatmap showing members of the “cholinergic synapse” gene set from KEGG. Different isoforms are marked by .1 and .2 next to the gene names.

In agreement with our scATAC-seq data (**Figure 4B-C and Figure S7C**), we found JUNB protein to be significantly upregulated in the brain of *Ppp1r15b*^R658C^ mice (**Figure 5A**). Although FOSL2 function correlated with persistent ISR activity in glutamatergic neurons (**Figure S7D**), no significant changes were observed in FOSL2 protein levels. Given the similarity in DNA-binding consensus sequences between these AP-1 transcription factors^50^ (**Figure S8**), we speculate that the chromatin opening around FOSL2 motifs may be attributed to JUNB. Interestingly, due to their known involvement in memory processes^51^, we observed reduced levels of several muscarinic acetylcholine receptors (mAChRs), including CHRM1, CHRM2, CHRM3, and CHRM4, as well as metabolic enzymes such as acetylcholinesterase (ACHE), which are critical for cholinergic synapse function, in *Ppp1r15b*^R658C^ mice (**Figure 5F**).

Lastly, we implemented a comprehensive integrative approach that has the potential to elucidate new mechanisms by which persistent ISR activity leads to cognitive dysfunction, which would significantly advance our understanding of its molecular underpinnings. Using SignalingProfiler^52^, a cutting-edge pipeline that integrates multi-modal data with prior knowledge and derives optimal coherent causal networks (**Figure 6A**), we analyzed ISR-driven signaling. Specifically, we combined three data sets—cell-type specific RNA-seq with bulk proteomics and phosphoproteomics—to construct initial networks representing glutamatergic and GABAergic neuronal contexts. The networks were then optimized using the CARNIVAL algorithm^53^, which identifies up-stream regulatory signaling, resulting in causal networks for GABAregic and glutamatergic neurons, respectively. As expected, ATF4 emerged as a central node in both context (red symbols; **Figure 6B-C**). Among the many intriguing candidates identified, JUNB stood out due to its selective regulation in glutamatergic neurons (**Figure 4**). More interestingly, this integrative and optimized approach predicts a potential cell type-specific mechanism of JUNB regulation. In GABAergic neurons, ITCH targets JUNB for degradation via ubiquitination^54^, thereby suppressing its activity (**Figure 6C**). In contrast, in glutamatergic neurons, JUNB is regulated by glycogen synthase kinase 3β (GSK-3 β), which typically suppresses JUNB activity through phosphorylation^55^. Given that GSK3β activity is predicted to be reduced in glutamatergic neurons, this inhibition is alleviated, allowing JUNB to become more active and induce its downstream transcriptional program (**Figure 6B**). These findings may explain the selective activation of JUNB (AP-1) in glutamatergic neurons under persistent ISR conditions. Furthermore, they underscore the utility of combining multi-modal datasets with advanced network-based approaches to uncover cell-type-specific regulatory mechanisms and generate new hypotheses. Finally, future studies will focus on functionally characterizing this pathway to deepen our understanding of its role in ISR-associated processes.

**Figure 6.**
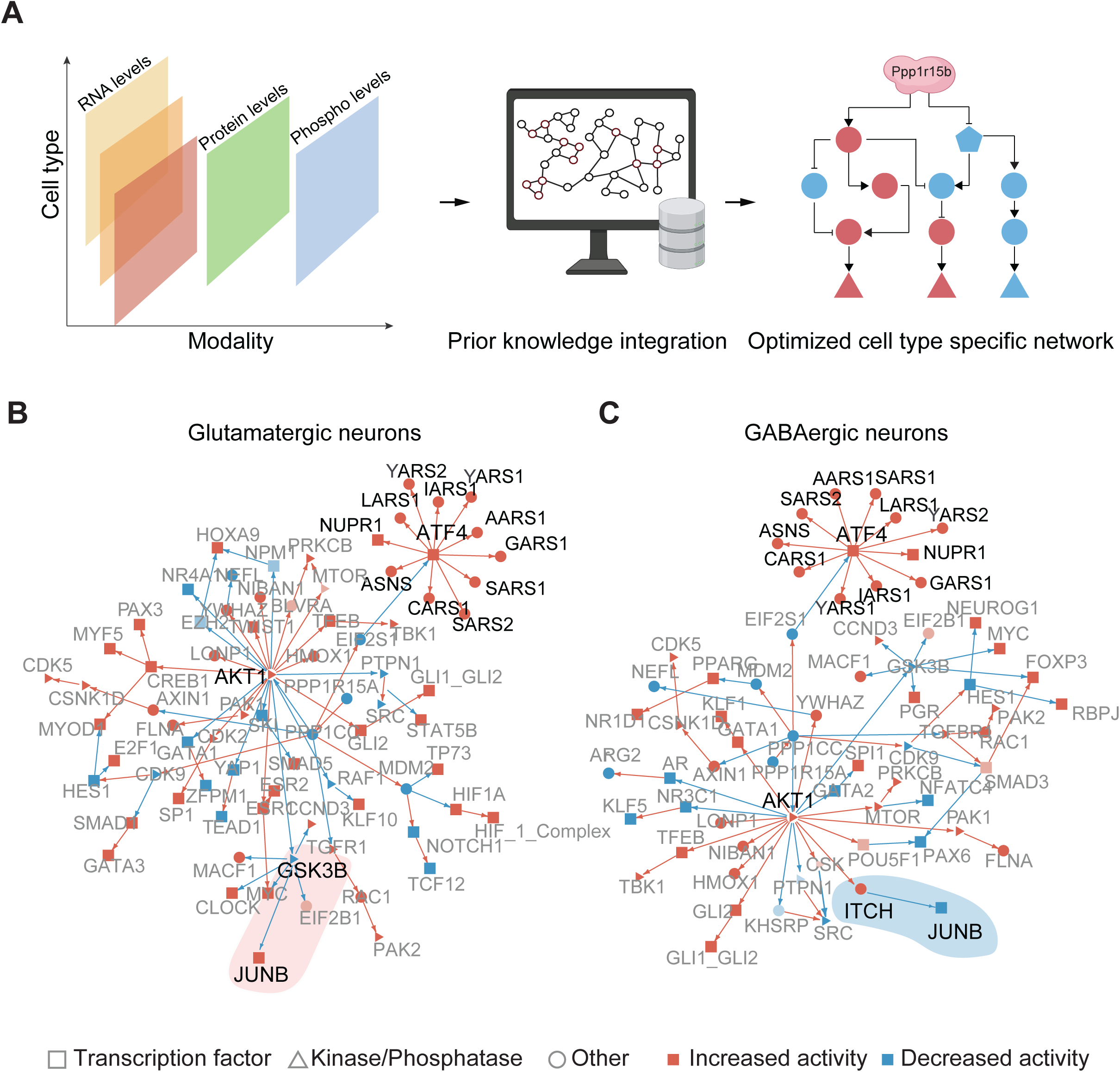
Multi-modal characterization of persistent ISR regulation. (**A**) Schematic illustrating the framework for integration of multi-omic data modalities from brain samples of WT and *Ppp1r15b*^R658C^ mice. Single-cell data (pseudo bulk counts), bulk proteomic and phosphoproteomic data are integrated with prior knowledge graphs. Cell type-specific networks are optimized, with PP1 as the input perturbation node, to identify the smallest coherent graphs that align with experimental data. (**B-C**) Functional protein network analysis for glutamatergic and GABAergic neurons in *Ppp1r15b^R658C^* brain samples, performed using SignalingProfiler. Nodes represent proteins, with red nodes indicating upregulated proteins and blue nodes indicating downregulated proteins. The edges between nodes represent protein interactions, with red lines denoting activating interactions and blue lines representing deactivating interactions, based on prior knowledge from SignalingProfiler’s annotated graph. Node shapes correspond to functional categories: triangles represent kinases, squares represent transcription factors, and circles denote other functional categories.

## Discussion

In this study, we present a comprehensive, multi-modal, cell-type-specific resource derived from the *Ppp1r15b*^R658C^ mouse model (see accompanying manuscript, Reineke *et al*.). This model provides a robust platform for dissecting the molecular mechanisms underlying ISR-mediated cognitive decline. First, it enables persistent ISR activation in a physiologically relevant context, without the confounding effects of secondary pathways disruptions commonly seen in other ISR-related models, such as those associated with Alzheimer’s disease^56^ and Down syndrome^57^. Second, the *Ppp1r15b*^R658C^ variant, which activates the ISR, directly leads to cognitive and synaptic plasticity deficits in the mutant mice, closely mirroring the human condition of intellectual disability. Most importantly, ISR suppression fully reversed the cognitive decline and synaptic dysfunction in this model (see accompanying manuscript, Reineke *et al.*), demonstrating that persistent ISR activation is the primary driver of the pathology.

By integrating multi-modal datasets—including scRNA-seq, scATAC-seq, proteomics, and phospho-proteomics—we generated an ISR atlas of the brain and multi-omic fingerprint of persistent ISR activation. This resource serves as a framework for generating and testing new hypotheses and advancing our understanding of how ISR activation contributes to cognitive decline. Specifically, it has allowed us to begin to address some critical questions, including how different brain cell types respond to ISR activation, the downstream effectors involved in ISR-mediated cognitive decline, and the potential to identify an ISR-driven molecular signature of cognitive dysfunction across neurological disorders.

A significant contribution of this study is the discovery that neurons and glial cells—e.g., astrocytes, oligodendrocytes, and microglia—respond to ISR activation in fundamentally different ways. These findings challenge the notion of a uniform ISR response across all brain cell types, highlighting the complexity of how dysfunctional stress pathways contribute to pathological outcomes in neurological disorders. Single-cell-omics revealed cell type-specific gene expression patterns in mice with persistent ISR activation (**Figure 1-2**). For instance, specific neuronal populations, such as GABAergic and glutamatergic neurons, showed a more prominent activation of the ISR compared to other cell types (**Figure 2E**). Moreover, single-cell recordings show that GABAergic, but not glutamatergic, synaptic transmission was increased in *Ppp1r15b*^R658C^ mice (see accompanying manuscript, Reineke *et al.*), indicating differential vulnerabilities to ISR activation. The increased GABAergic synaptic function in *Ppp1r15b*^R658C^ mice likely impairs cognitive function by disrupting the excitatory-inhibitory balance, which is critical for proper cognitive function^58^. Several lines of evidence support this hypothesis. Genetic inhibition of the ISR reduces GABAergic synaptic transmission without affecting glutamatergic synaptic transmission in normal mice^7^ and models of persistent ISR activation^21^ (see accompanying manuscript, Reineke *et al.*). Increased GABA inhibition impairs both LTP and long-term memory formation^59^. Finally, inhibition of the ISR in a subtype of GABAergic neurons facilitates long-term memory formation^12^.

Recent molecular genetic studies show that genetic inhibition of the ISR in other neuronal subtypes, such as glutamatergic or cholinergic neurons, also facilitated long-term memory formation^12,60,61^. Partial genetic inhibition of the ISR also promoted LTP in dopaminergic neurons and enhanced drug-induced behaviors^62,63^. However, complete blockage of ISR signaling in dopaminergic neurons impairs long-term spatial and aversive associative memories^64^, suggesting that baseline ISR activity in these cells is essential for proper mnemonic processing. Since aberrant cholinergic function is linked to age-related cognitive impairment and Alzheimer’s disease^51,65^, this pathway may represent an additional mechanism through which ISR activation impacts cognition.

Another important finding in our study is the identification of distinct ISR effectors in different neuronal subtypes (**Figure 4**). JUNB, an AP-1 transcription factor, mediated the ISR response in glutamatergic neurons, while ATF4 dominated in GABAergic neurons. The results from our multi-omic integrative approach (**Figure 6A**) predicts that JUNB is degraded in GABAergic neurons through the activity of the E3 ligase ITCH, which is absent in glutamatergic neurons during persistent ISR activation (**Figure 6C-D**). A recent study has linked AP-1 signaling to aging and disruption of youthful cell identity^66^, raising the possibility that ISR-mediated modulation of AP-1 may influence neuronal aging or rejuvenation. Future studies will need to explore the link between the ISR and AP-1 in glutamatergic neurons and the potential aging of specific brain cell populations by persistent ISR activation. More importantly, our findings also revealed that *Atf4* deletion in GABAergic neurons—but not in glutamatergic neurons—impacts ISR-mediated cognitive decline, functionally demonstrating that different neuronal subtypes rely on unique ISR effectors to regulate long-term memory. These results emphasize the importance of studying persistent ISR activation within specific cell populations, as the molecular players involved in ISR-induced cognitive dysfunction vary across cell types.

In contrast to the *Ppp1r15b*^R658C^ model, a Vanishing White Matter model, characterized by ISR activation due to a loss-of-function variant in eIF2B (*Eif2b5*^R191^ mice)^67^, shows that ISR activation primarily affects astrocytes and oligodendrocytes^67^, whereas in the *Ppp1r15b*^R658C^ model, it mainly impacts neurons. Neurons, which depend on precise protein synthesis regulation for synaptic plasticity and memory^68–71^, are likely more vulnerable to ISR-induced cognitive decline, while defects in glial cells could lead to impaired myelination and neuroinflammation, key features of vanishing white matter disease. Indeed, it will be interesting to examine whether neurons are the most susceptible population in other human cognitive disorders (or mouse models) carrying ISR-activating variants, such as those linked to the MEHMO (mental deficiency, epilepsy, hypogenitalism, microcephaly, and obesity) syndrome, a genetic ISR-related and X-linked disorder^18–20,72^. These findings, which underscore the importance of studying cell-type-specific ISR responses, suggest that the pathological consequences of persistent ISR activation depend on the cell type involved.

Finally, our single-cell-omic studies identified a molecular signature of persistent ISR activation, which can serve as a biomarker for cognitive dysfunction. We defined a set of ISR-regulated genes and pathways that consistently appear across different models of cognitive decline, including neurodevelopmental disorders and neurodegenerative diseases, as well as age-related cognitive decline (**Figure 3**). This comprehensive ISR signature extends beyond ATF4 targets and allows the molecular assessment of ISR activation across tissues or disease states.

In summary, our findings provide a novel perspective into the mechanisms behind persistent ISR activation and its role in cognitive decline. Furthermore, the comprehensive multi-omic framework we have developed opens new avenues for studying ISR biology in the brain, which promises to accelerate the development of innovative approaches for diagnosing and treating cognitive disorders of different etiologies.

## Methods

### Nuclei isolation from mouse brain tissues

Nuclei isolation was performed following a previously established protocol^73^ with the following modifications. Approximately 5-10 mg of frozen cortical tissue was homogenized in 500μL of nuclei extraction buffer [0.32 M sucrose, 10mM Tris pH 7.4, 5mM CaCl_2_, 3mM Mg acetate, 1mM DTT, 0.1 mM EDTA, 0.1% Triton X-100, 0.2U/ml Protector RNAse inhibitor (Sigma cat. 3335399001)] using a dounce homogenizer. The homogenized sample was then filtered through a 70mm filter. The sample was gently layered over a 750μl sucrose solution (1.8 M sucrose, 10 mM Trish pH 7.4, 3mM Mg acetate, 1 mM DTT) and then centrifuged at >16,000 x g for 30 minutes at 4°C. Nuclei permeabilization was conducted as per the 10X Genomics protocol (CG000375 Rev B; https://www.10xgenomics.com/support/epi-multiome/documentation/steps/sample-prep/nuclei-isolation-from-complex-tissues-for-single-cell-multiome-atac-plus-gene-expression-sequencing). Briefly, nuclei were resuspended in 100μl lysis buffer (10mM Tris-HCl pH 7.4, 10 mM NaCl, 3mM MgCl_2_, 1% BSA, 0.01% Tween 20, 0.01% NP-40, 0.001% digitonin, 1 mM DTT, 1U/mL of Protector RNAse inhibitor) and incubated for exactly 2 minutes on ice. Nuclei were then gently washed one time prior to being resuspended in 30mL of 1X nuclei buffer with 1 mM DTT and 0.5U/μl of Protector RNAse inhibitor. Nuclei quality and concentrations were then determined using the Countess II FL.

**Single nucleus multi-omics.** Transposition, nuclei isolation, barcoding, and library preparation were performed following the 10X Genomics Chromium Next GEM Single Cell Multiome protocol (CG000338 Rev F; https://www.10xgenomics.com/support/epi-multiome/documentation/steps/library-prep/chromium-next-gem-single-cell-multiome-atac-plus-gene-expression-reagent-kits-user-guide), with the following alterations. Samples were processed in two batches (*n* = 14), evenly divided by genotype and sex to ensure efficient handling while maintaining sample integrity. Each sample was loaded onto a single lane of the Chromium Next GEM Chip J according to the manufacturer’s recommendations to recover approximately 10,000 nuclei per lane. Sequencing libraries were prepared and subsequently sequenced by Novogene using Illumina NovaSeq X.

### Single-cell multi-ome data pre-processing and analysis

Raw sequencing reads were aligned to the GRCm38 mouse reference genome using the Cell Ranger ARC pipeline (v2.0.2). The resulting raw RNA count matrix and ATAC fragment data were further processed using R packages Seurat^74^ (v5.2.0) and Signac^30,31^ (v1.14.9), respectively. ATAC reads were re-quantified based on the union of consensus accessible chromatin regions (ACRs) between the samples. These ACRs were filtered for to include regions with peak width between 20 and 10,000 base pairs, after which the samples were merged for subsequent analysis. Subsequently, RNA quality filtering was performed using PopsicleR^75^ (v0.2.1), applying strict thresholds to ensure high data quality. The criteria for filtering included: the maximum number of genes detected per cell (G_RNA_low=500 and G_RNA_hi=Inf), the range of molecules detected per cell (U_RNA_low=-Inf and U_RNA_hi=37000), and the upper limits for mitochondrial, ribosomal, and dissociation gene percentages (percent_mt_hi=10, percent_ribo_hi=100, and percent_disso_hi=100, respectively). We then filtered based on ATAC metrics (500 < nCount_ATAC < 100,000, nucleosome_signal < 3, TSS.enrichment > 1). After these combined filtering steps, 85,402 cells met the minimal inclusion for analysis.

The data set exhibited an average transcriptomic read depth is 6,919 unique reads per cell and 10,012 unique ATAC reads per cell. On average, 2,598 unique genes and 8,602 unique chromatin peaks were detected per cell. RNA normalization was performed using the log-normalization method within the Seurat’s NormalizeData function^74^, followed by scaling (ScaleData). Principal component analysis (PCA) was then conducted on the top 2,000 highly variable features, retaining the first 30 principal components for downstream analysis.

For ATAC data, latent semantic indexing (LSI, dimensions 2:30) was used for dimensionality reduction, which involved applying RunTFIDF, FindTopFeatures, and RunSVD functions from Singac^30,31^, focusing on chromatin feature matrices and retaining dimensions 2 through 30. Integration of RNA and ATAC modalities was achieved by computing a weighted neighborhood graph that leveraged both datasets. Finally, Uniform Manifold Approximation and Projection (UMAP) was used to project the integrated data into a two-dimensional space for visualization.

### Dataset Annotation

We annotated the datasets using a dual-pronged approach to ensure comprehensive and accurate cell type identification. First, we utilized the MakeAnnotation function within PopsicleR ^75^, which integrates two reference-based annotation tools: SingleR^76^ and scMCA^77^. SingleR leverages bulk RNA datasets, specifically mouse RNA-seq, as its reference while scMCA uses single-cell Mouse Cell Atlas (scMCA) to annotate cell populations. In parallel, we employed a reference mapping approach implemented in Seurat^74^. This method identifies “anchors” between the low-dimensional structures of reference and query datasets, enabling the transfer of cell annotations from the reference to the query dataset. For this process, we downloaded the Allen Brain Atlas Mouse whole-brain transcriptomic cell type atlas^32^ (Smart-seq v4 dataset) as the reference. Using Seurat’s FindTransferAnchors function, we performed unsupervised anchoring based on the first 30 principal components of both the reference and query datasets.

After completing both the reference mapping and automated annotation processes, we merged the results to obtain a unified annotation scheme. For downstream analyses, we categorized the dataset into three annotation levels. Level 1 distinguished between GABAergic neurons, Glutamatergic neurons, and non-neuronal populations. Level 2 provided finer resolution by further subclassifying non-neuronal populations. Finally, Level 3 offered the highest granularity by subdividing neuronal populations into specific subtypes.

### Identifying differentially abundant and differentially expressed subpopulations

We applied the MiloDE and SEAcells frameworks for differential abundance and expression. The MiloDE framework assigns cells to partially overlapping neighborhoods on a k-nearest neighbor (kNN graph) rather than using discrete clusters, allowing for more nuance differential abundance and differential expression analyses. For differential abundance testing, we used the MiloR package (v1.1.0), which accounts for the overlap between neighborhoods using a weighted Benjamini-Hochberg correction, where the *P*-values are weighted by the reciprocal of the connectivity between neighborhoods. For differential expression analyses, the MiloDE package (v0.9) builds on this framework, introducing modifications to neighborhood assignment, statistical testing, and multiple testing correction. Specifically, MiloDE uses second order kNN graphs to better capture the local structure of the subpopulations, and the differential expression analysis is performed using edgeR. This framework also incorporates corrections for connectivity and neighborhood size. In our analyses, neighborhood graphs were constructed using parameters k=25, d=30 in PCA space. For differential expression, we ran edgeR with the covariates of sex and genotype, and design matrix of design = ∼ sex + geno. Genes were considered significantly differentially expressed if they satisfied two conditions: *P*-value corrected across genes < 0.05 and *P*-value corrected across neighborhoods < 0.05.

In addition to MiloDE, we employed the SEAcells^39^ framework, which enables the aggregation of single cell of cell states into metacells while preserving biological heterogeneity. Metacells provide a robust way to detect subtle shifts in granular cell states, even when fold changes are small. SEAcells operates by leveraging a graph-based manifold learning approach built on the nearest-neighbor graph of the dataset. To do this, waypoint cells from the neighborhood graph are first selected using a minimum-maximum sampling approach strategy. These waypoint cells serve as anchors for subsequent iterative assignments of individual cells into metacells, based on archetypal analysis. This transformation group cells into compact clusters under strict similarity conditions while preserving outlier states in the phenotypic space. For our dataset, we constructed the SEAcell kernel transcriptomic PCA space using the first 10 principal components as initialization. Each metacell was designed to aggregate approximately 75 cells. The algorithm was run for 50 iterations and reads from WT and *Ppp1r15b*^R658C^ cells in each metacell were summarized using their soft assignment. Afterwards, the resuting metacell data was normalized, scaled, and processed for PCAs using the Seurat pipeline. We then performed differential expression analyses using Seurat’s FindMarkers function within each level 3 (l3) cell type, with the following parameters: min.pct = 0, logfc.threshold = 0.15,min.cells.feature = 0,min.cells.group = 0. Finally, we intersected the differentially expressed genes identified in both the MiloDE and SEAcells analyses.

### Gene module identification

We performed scWGCNA^78^ on the identified miloDE neighborhoods to identify co-expression modules. Briefly, scWGCNA is the single-cell adaptation of Weighted Gene Network Co-Expression Analysis (WGCNA), where modules of co-expressed genes are identified based on their expression patterns across metacells. Matrices for topological overlap and adjacency, are calculated, with genes being assigned to discrete co-expression modules. Initial gene memberships are pruned based on the requirement of correlation with the overall expression of their co-expression module. Given the strong dependence of module size and the number of modules on the number of neighborhoods a gene must be significant across (defined by n_hoods_sig.thresh), we iterated through values ranging from 1 to 58. We finally settled on paramaters (n_hoods_sig.thresh = 8, npcs = 10, pval.thresh = 0.1) as the most representative of co-expression modules in our dataset. Through this process, we identified two modules: one module containing genes with uniformly increasing expression and another module with decreasing expression, both appearing consistently across iterations. Additionally, we identified a module specific to oligodendrocytes, which was present in fewer than 50% of the iterations. Finally, we intersected these co-expression modules with the list of significant genes in both SeaCells and MiloDE to derive our final ISR *up* (28 genes) and ISR *down* (12 genes) signatures.

### Gene Enrichment Analysis

Gene enrichment analyses were performed using two tools: the web-based tool, Metascape^79^, and GSEA using the package WebGestaltR (doi: 10.1093/nar/gkz401, v0.4.5). For Metascape, we provided our gene sets as input and all detected genes as the background. Significant Gene Ontology (GO) terms were selected with the threshold of adjusted *P*-value < 0.05. For webGestaltR, we input differentially expressed genes between WT and *Ppp1r15b*^R658C^ cells (adjusted *P*-value < 0.05) for each of the cell types. Genes were ranked according to ± (avg FC) X -log10 (adj *P*-value). We used the following pathways for enrichment analyses: KEGG pathway, Gene Ontology (Biological Process, non-redundant) and Gene Ontology (Molecular Function, non-redundant for each l2 level cell annotation.

### Validation in external datasets

We obtained publicly available datasets for validation, including those related to human Down Syndrome^40^ (EGAS00001005691), Parkinson’s disease and Lewy Body Dementia^42^ (GSE178265) and aged mouse data^80^ (GSE207848), either through by direct download or from the CZ CELLxGENE CENSUS^81^. For validation, we created gene sets of orthologous genes of our up and down gene sets. We then used UCell^82^ to calculate signature scores based on the Mann-Whitney U statistic. We then performed Kruskal-Wallis tests (KW) tests between WT and *Ppp1r15b*^R658C^ cells within each l2 level cell annotation using the stat_compare_means function. Our upregulated and downregulated gene sets were validated in mouse samples, and the ISR signature in the human disease datasets. As a final step of validation, we calculated a single-cell disease relevance score (scDRS) for each metacell^43^. This method links scRNA-seq data with polygenic disease risk, in a cell type independent mode. We used the pre-calculated disease/magma scores for mouse brain data shown exhibited in the example vignette.

### Downstream single-cell epigenetic analysis

We identified ACRs within all samples using MACS3^83^ followed by filtering to standard chromosomes, and then merging the accessible regions across individual samples. For downstream analyses, we created fragment files based on their SEAcells metacell assignments. We then computed gene activities, and the deviations in accessibility of transcription factors using the GeneActivity, and RunChromVAR functions in Signac^30,31^. We obtained position weight matrices from the JASPAR2020^84^ database for vertebrates. Motif positioning in open regions of chromatin was determined via the Signac function FindMotifs, and the R package motifmatchr (v.1.24.0). Significant transcription factors (TFs) were identified using FindMarkers (adjusted *P*-value < 0.05, test.use = ’LR’, latent.vars = ’nCount_peaks’). We then linked accessible regions to normalized expression changes (using SCTransform) for all genes using the LinkPeaks Signac command (*P-*value < 0.001 & z-score > 0). To assess transcription factor footprinting in these accessible regions, we used the Footprint function within Signac. Additionally, to explore cell type-specific epigenetic changes linked to the ISR, we created a filtered list of regulatory elements. ACRs were classified as linked to the ISR if they met the following criteria: 1) they were differentially accessible between WT and *Ppp1r15b*^R658C^ cells within each l3 cell annotation (*P*-value < 0.005), 2) they were significantly linked to a gene ± 50 kbps from the TSS, (*P*-value < 0.001 & z-score > 0), and 3) the corresponding gene was differentially expressed between WT and *Ppp1r15b*^R658C^ cells within the same L3 cell annotation (adjusted *P*-value < 0.05). Finally, the results were aggregated the on their l1 cell annotation, revealing 134 opening and 57 closing regulatory elements. Hpergeometric tests were performed using the FindMotifs function (*P*-value < 0.05), and peaks were annotated using ChIPseeker^85^ (v1.42.0).

### Building gene regulatory networks

We used a gene regulatory network (GRN) inference tool Pando^46^ (v1.1.1) to identify TF activity in a cell type-specific manner. Similar to our in-house approach described above, Pando integrates both ATAC and RNA modalities to simultaneously infer regulatory elements for each gene. Pando models expression changes of genes via: 1) selecting linked regulatory genes within ± 10 kbps from the transcription start site (TSS), 2) scanning these regions for TF motifs (for this we used the custom motif point weight matrices provided by Pando), 3) identifying TF-region pairs for each gene, and 4) constructing regression models where the region-TF pairs serves as independent variables and the target genes serve as dependent variables. We applied Pando to our 28 upregulated ISR signature genes, and then to all 163 differentially expressed genes that were identified in both MiloDE, and SEAcells. For this analysis, we focused on TFs that were differentially expressed between WT and *Ppp1r15b*^R658C^ cells within l3 level cell annotations. We used the find_modules command with the following parameters: p_thresh = 0.1, nvar_thresh = 2, min_genes_per_module = 1, rsq_thresh = 0.05 to identify co-regulated modules. Finally, we calculated cell-type-activity of a TF by multiplying the mean regulatory coefficient (coef) with the average expression of the TF within each cell type.

### Mass Spectrometry Sample preparation

The cortex from mice was homogenized in 100 mM EPPS buffer (pH 8.5) containing 1 % sodium deoxycholate and heated to 95 °C for 5 minutes. Proteins (500 µg) were reduced by addition of 1 mM DTT and heated at 25 °C for 15 minutes, then alkylated with 5 mM iodoacetamide at 25°C for 15 min in the dark. Samples were digested with Lys-C (1:50 enzyme-to-protein-ratio) for 6 hours at 25 °C (1000 RPM), followed by trypsin (1:50 enzyme-to-protein ratio) for 16 hours. Peptides were labeled with TMT dissolved in dry acetonitrile. Following a label-check by mass spectrometry, samples were quenched with 8ul of 1M Tris and pooled. The combined material was acidified (pH < 2) with trifluoroacetic acid and desalted using SDB-XC (Phenomenex, Torrance, CA). Lyophilized peptides were separated by basic reverse-phase chromatography, as previously described^86^ with the following modification: mobile phases were buffered exclusively with 10 mM ammonium bicarbonate. Fractions 13-84 were resuspended in 80 % acetonitrile (ACN), 0.1 % trifluoroacetic acid (TFA) and concatenated every 24th fraction. Ninety percent of the peptide underwent phospho-enrichment using Fe-IMAC using (Agilent AssayMap Bravo protocol v.2.1), while the remaining 10% was diluted to 5% ACN, 0.1% TFA for full proteome analysis.

Full proteome samples were separated using a two-step linear gradient on a 25 cm Ionopticks Aurora Ultimate column maintained at 50 °C. Mobile phase A consisted of 0.1% formic acid, while mobile phase B contained 80% acetonitrile with 0.1% formic acid. The gradient progressed from 6% to 40% B over 120 minutes, followed by an increase to 50% B over 10 minutes.

Ions were generated using an Easyspray source with a spray voltage of 1500 V and an ion transfer tube temperature of 305 °C. A Thermo Fisher Scientific Orbitrap Ascend mass spectrometer was used to analyze TMT-labeled peptides with a fixed duty cycle of 2.5 seconds. MS1 scans were acquired over a 400–1600 m/z range using an Orbitrap analyzer at a resolution of 120,000.

For data-dependent acquisition, MS2 scans were performed in the linear ion trap in Turbo mode, using a maximum injection time of 35 ms. Peptides were isolated via quadrupole selection (0.7 m/z window) and fragmented with CID at 30% collision energy. A real-time search filtered peptides selected for MS3, allowing up to one missed cleavage and two methionine oxidations. The top 10 MS2 fragments were further fragmented with HCD at 55% collision energy and analyzed in the Orbitrap at a resolution of 45,000 over a 100–500 m/z range.

Phosphopeptides were analyzed using the same method as full proteome samples, with the following modifications: the spray voltage was increased to 1600 V. Data-dependent MS2 scans were performed following multi-stage activation, assuming a neutral loss mass of 98 Da, and measured in the Orbitrap at a resolution of 30,000. For MS3 selection, up to three variable modifications, two missed cleavages, and neutral loss masses of 79.97, and 97.98 were enabled for the real-time-search.

All .RAW files were processed in Proteome Discoverer v.3.0 against a mouse database from Uniprot (downloaded on March 16, 2023) and a common contaminants.fasta file from MaxQuant. The analysis followed the SPS-MS3 real-time-search template, incorporating Inferys rescoring for Sequest HT and Percolator for both Inferys and Comet search outputs. Reporter ion quantification was intensity-based, allowing 75% co-isolating, whit at least 65% of SPS mass matches corresponding to the precursor. Peptides with variable modifications were excluded from protein quantification.

### Processing and statistical analysis of proteomic data

Prior to statistical analysis, tandem mass tag (TMT) data were normalized by calculating channel-specific correction factors. These factors were determined by computing ratios between the maximum sum intensity and all other channels to correct for unequal sample loading. All normalization factors were below 1.2. Data were then filtered for missing values, and only proteins with at least 2 values per group (i.e. WT or *Ppp1r15b^R658C^*) were retained for statistical analysis resulting in a total of 9,621 proteins. Differential expression was tested using the ROTS function with 500 bootstraps and requiring K=500 for the number of differential hits^87^. Enrichment analysis was performed against the KEGG database using the R package WebGestaltR with the GSEA method ^88^. Only gene sets containing at least 5 members were tested. Categories enriched at < 5% FDR level are reported here.

### Multi-modal data integration with Signaling Profiler

We utilized SignalingProfiler to construct a contextualized coherent causal network by integrating multi-modal -omics data from this study with a large prior knowledge network (PKN). The PKN included curated protein interactions with known regulatory effects (e.g. activation or inhibition), documented phosphorylation events, and mapped regulatory relationships between kinases, transcription factors, and their targets. The major sources of information for the PKN included: PhosphoSitePlus and SIGNOR for phosphorylation events and kinase regulation, and OmniPath for transcription factor regulation and protein-protein interactions. W To model cell-type-specific signaling, we constructed two one-layer networks using maximal length of four nodes for GABAergic and glutamatergic neurons, respectively. Cell-type specificity was derived from pseudo-bulk sigle cell data, while proteomic and phosphoproteomic input data were the same for both networks. Protein phosphatase 1 (PP1) was added as a perturbation node to help with network initialization, given its role in ISR regulation. The networks were then optimized using the one-step CARNIVAL algorithm ^53^ and the cplex solver with default settings.

### Contextual Fear Conditioning

Experiments were conducted as previously described^21^. Briefly, mice were handled for 3 days (5-10 minutes each day) and acclimated to the conditioning chamber for 20 minutes for two additional days. On the training day, mice were placed in the chamber for 2 minutes (naïve phase), followed by two-foot shocks (0.75 mA, 2 sec, 90 sec apart). Afterward, the mice remained in the chamber for an extra minute before being returned to their home cages. Twenty-four hours later, mice were re-exposed to the same chamber for 5 minutes, and their “freezing” behavior (immobility except for respiration) was recorded using real-time video and analyzed with FreezeView software (Actimetrics, Lafayette, IN).

### Data sharing and availability

Using the ShinyMultiome.UiO package (https://www.biorxiv.org/content/10.1101/2023.06.20.545756v2), we developed an interactive web application that facilitates ISR-driven multi-omics data visualization, exploration, integration. This application can be accessed via the following link: https://altoslabs.shinyapps.io/ISR_atlas/.

## Supporting information

Supplementary Fgures and Legends

## Acknowledgements

This work was supported by Altos Labs, Inc. and NIMH (M.C.-M.). We thank members of the animal facility for support and members of the MCM and PW labs for comments on the manuscript.

