## Supplementary Fgures and Legends for "Mapping the ISR Landscape in Cognitive Disorders *via* single-cell multi-omics"

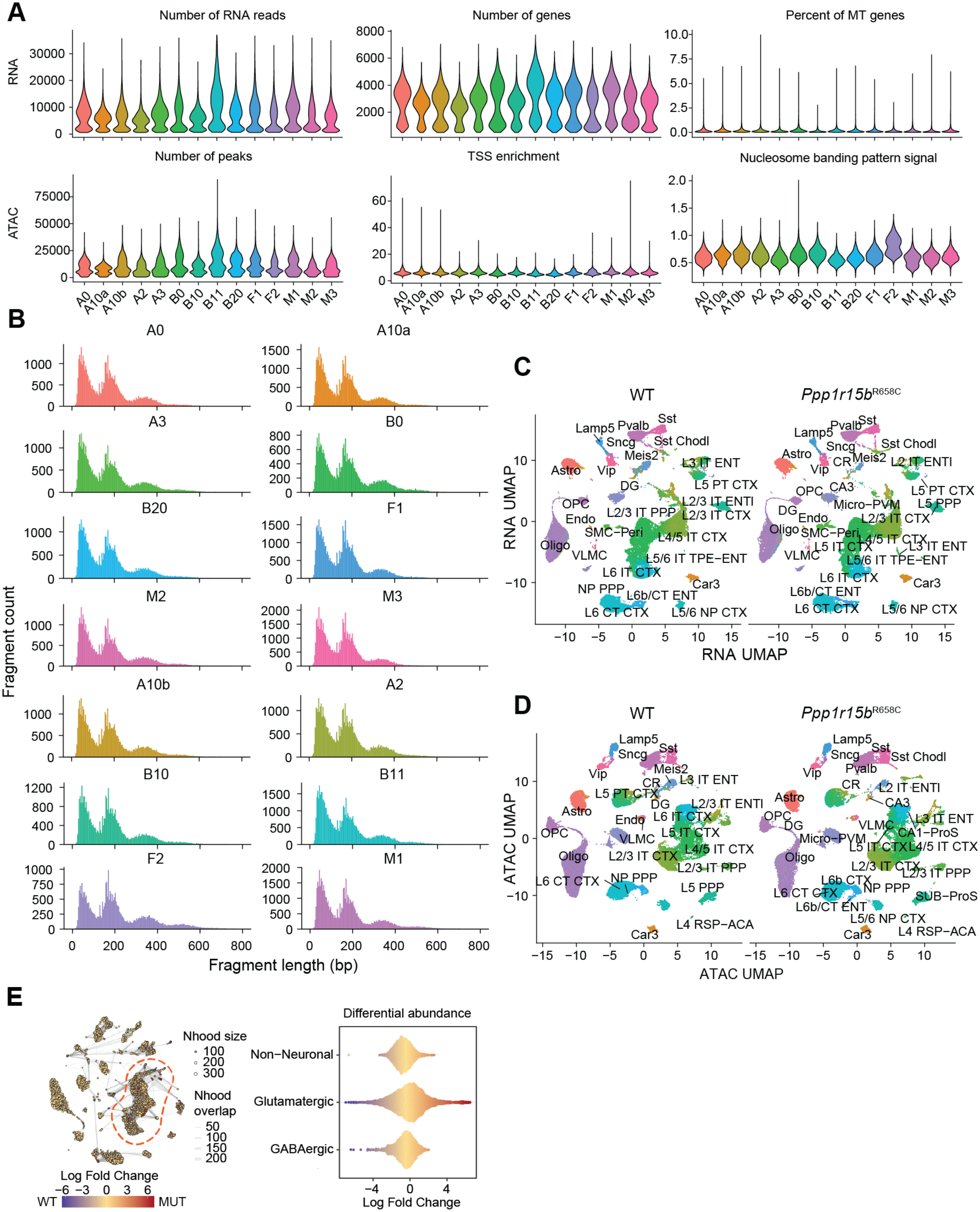


**Fig. S1.** Torkenczy *et al.*

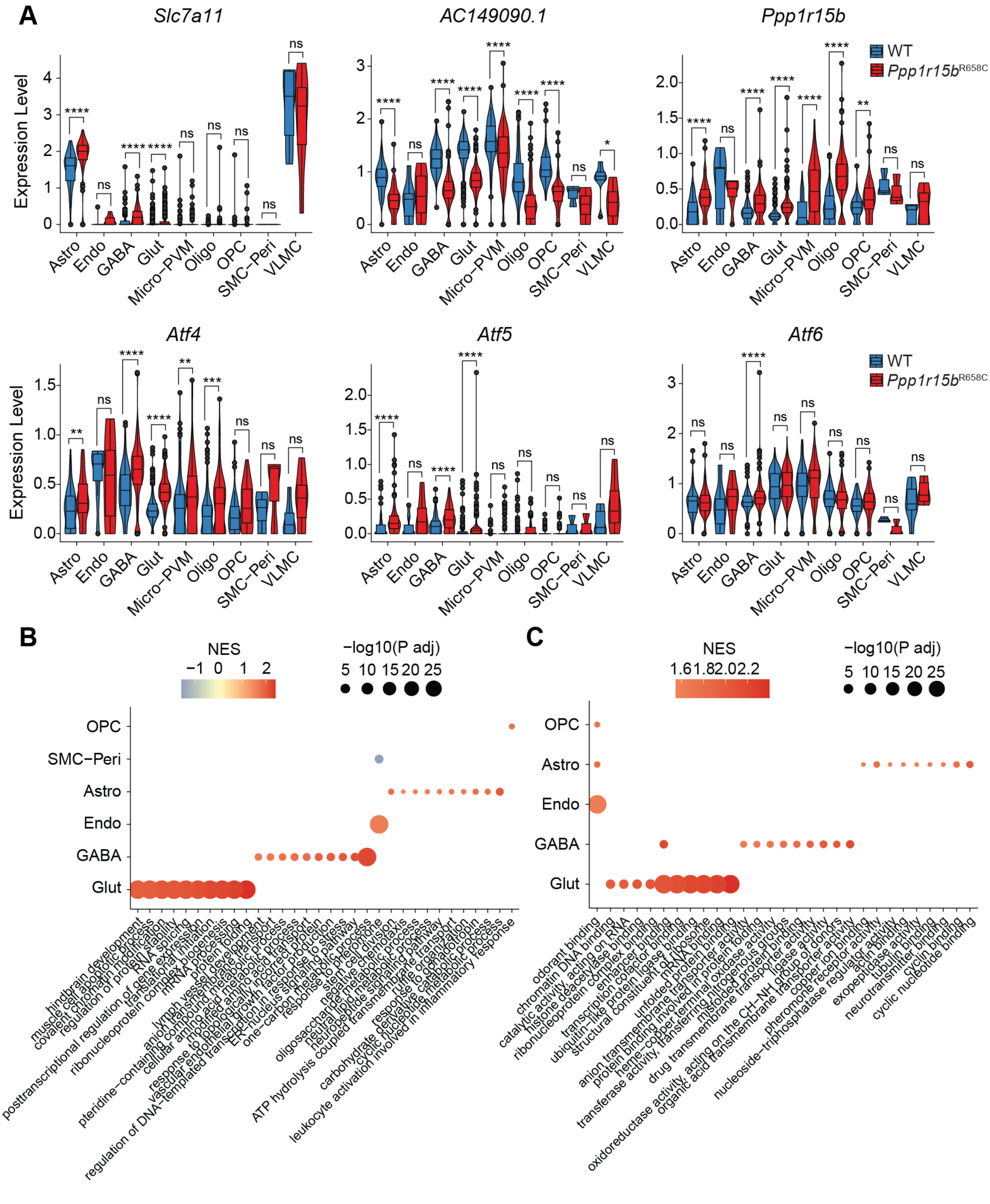

**Fig. S2.** Torkenczy *et al.*

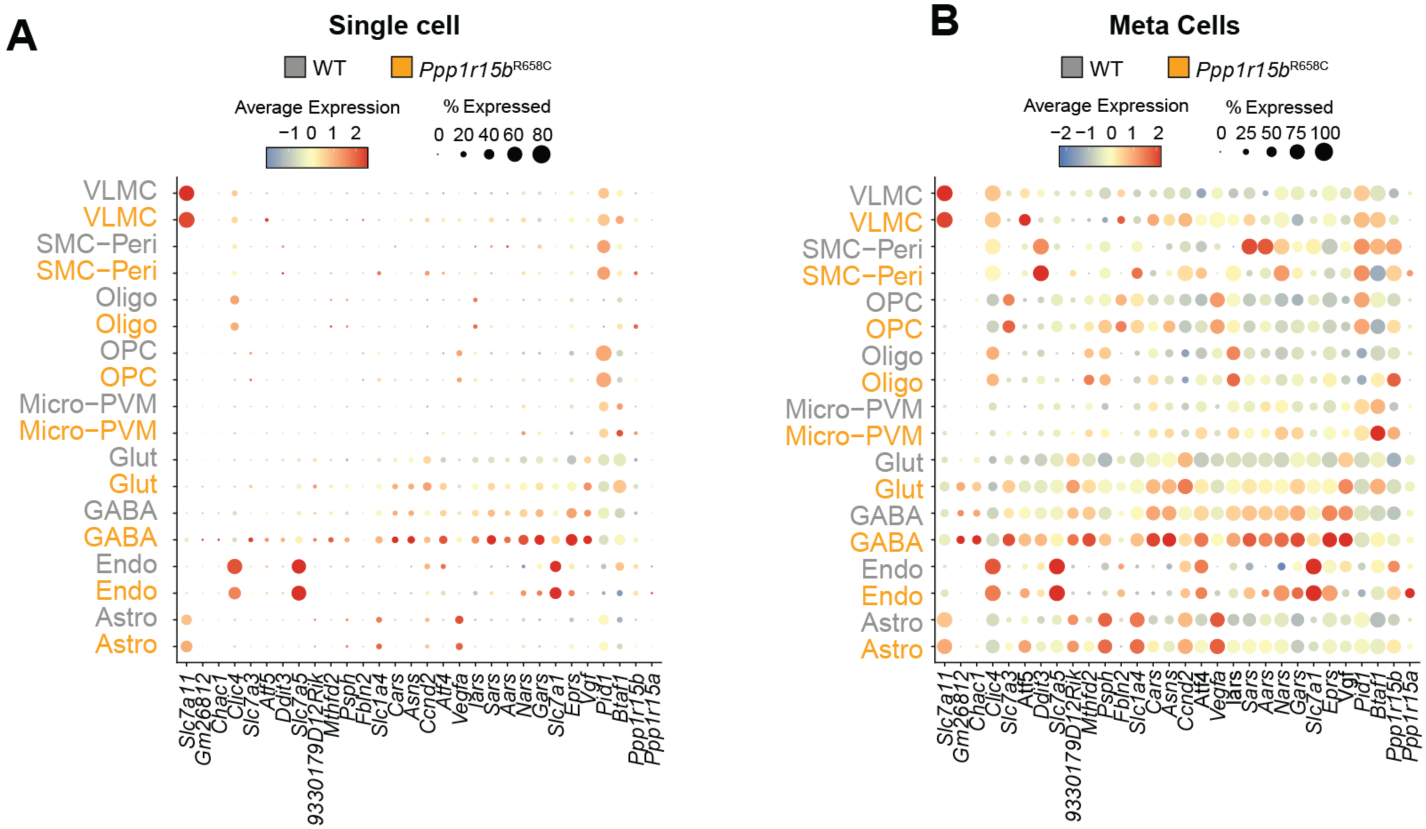


**Fig. S3.** Torkenczy *et al.*

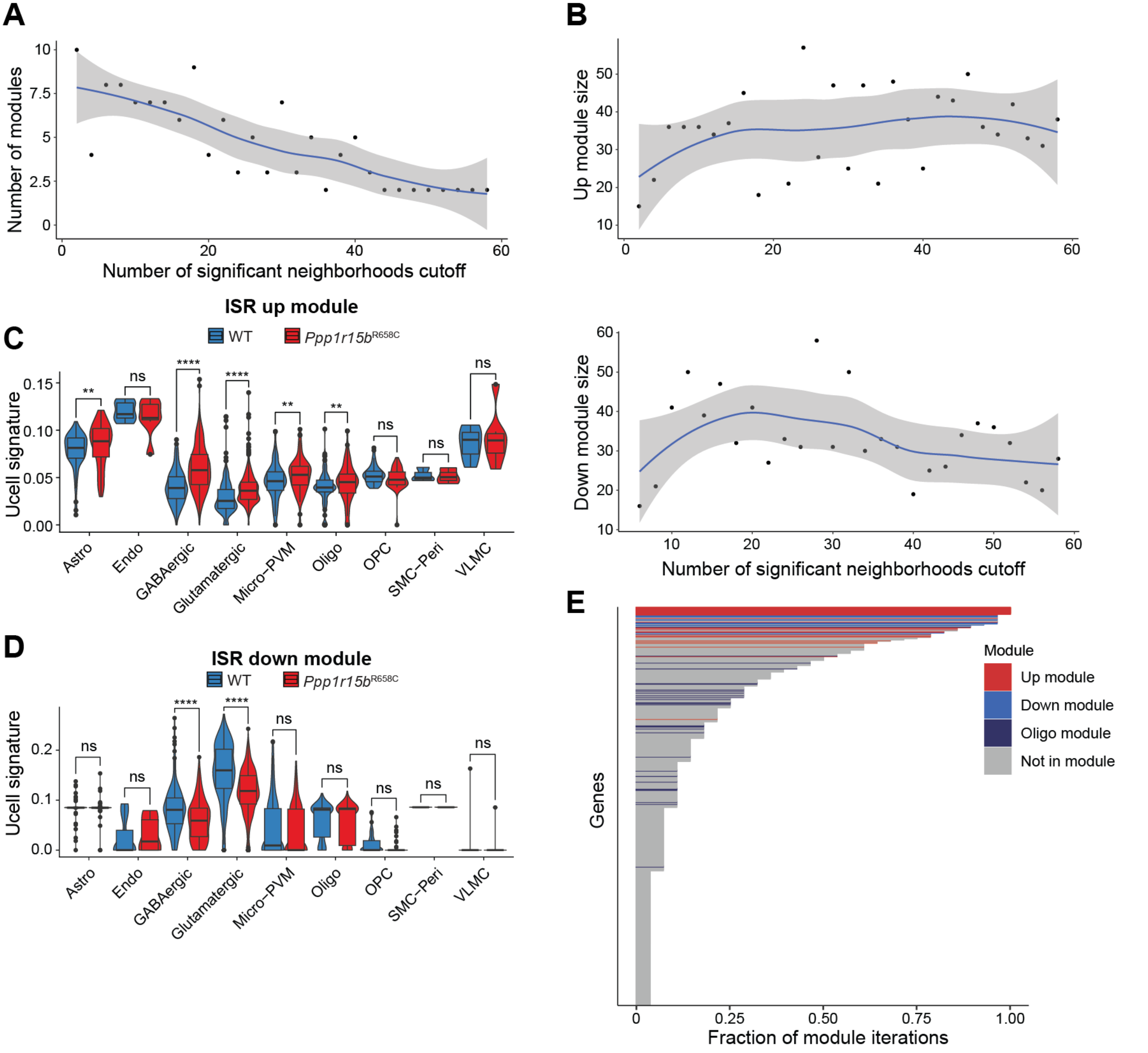


**Fig. S4.** Torkenczy *et al.*

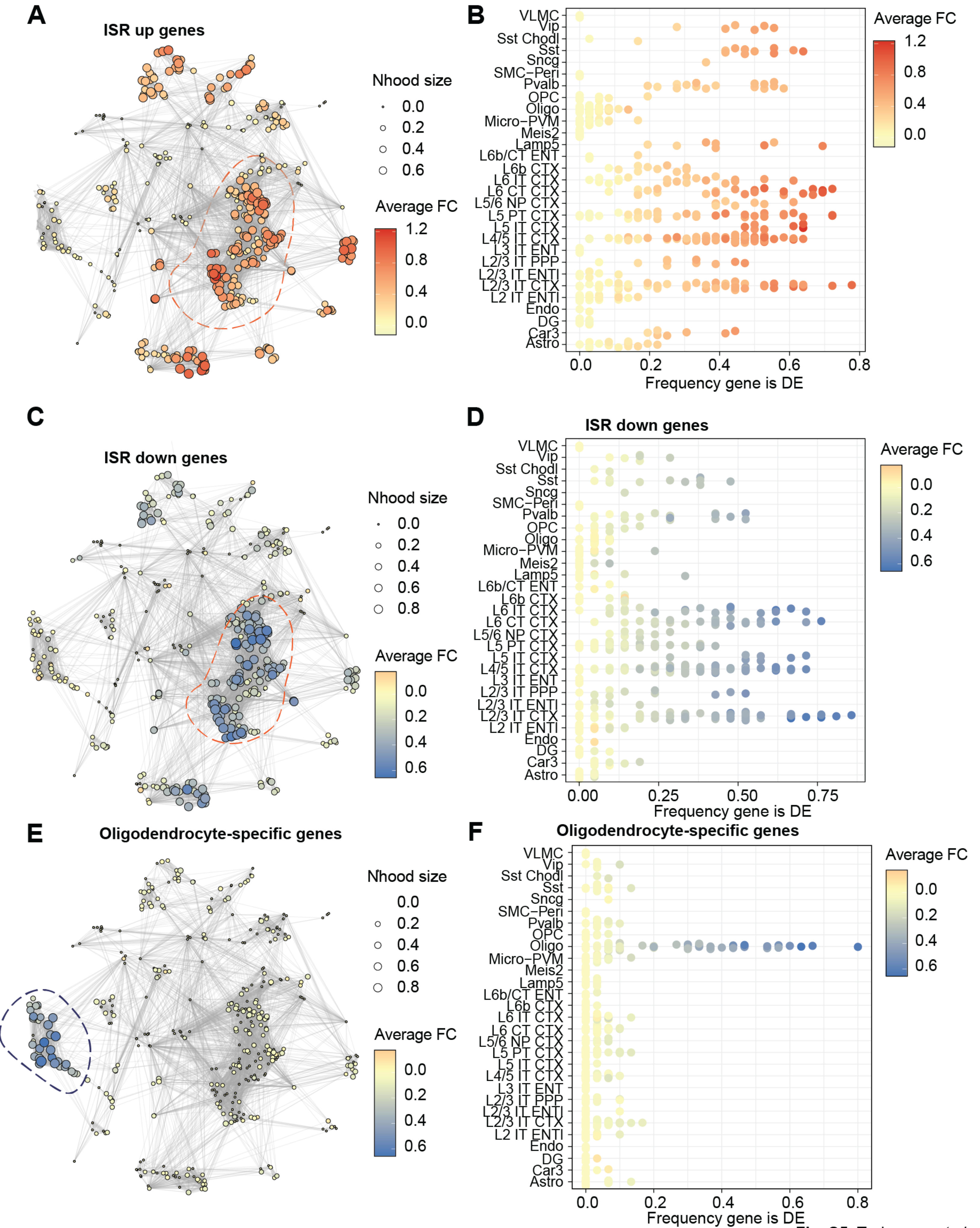


**Fig. S5.** Torkenczy *et al.*

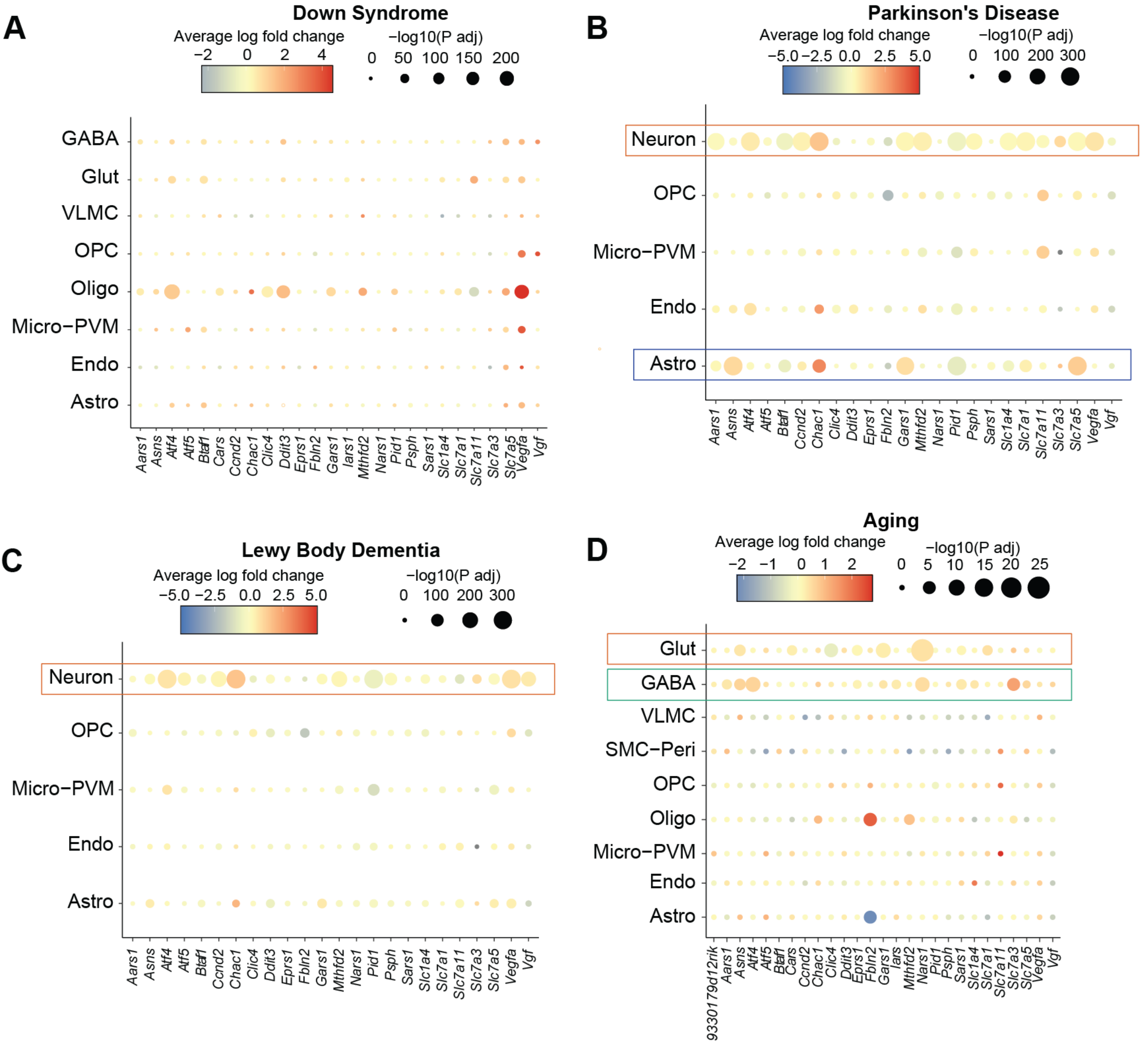


**Fig. S6.** Torkenczy *et al.*

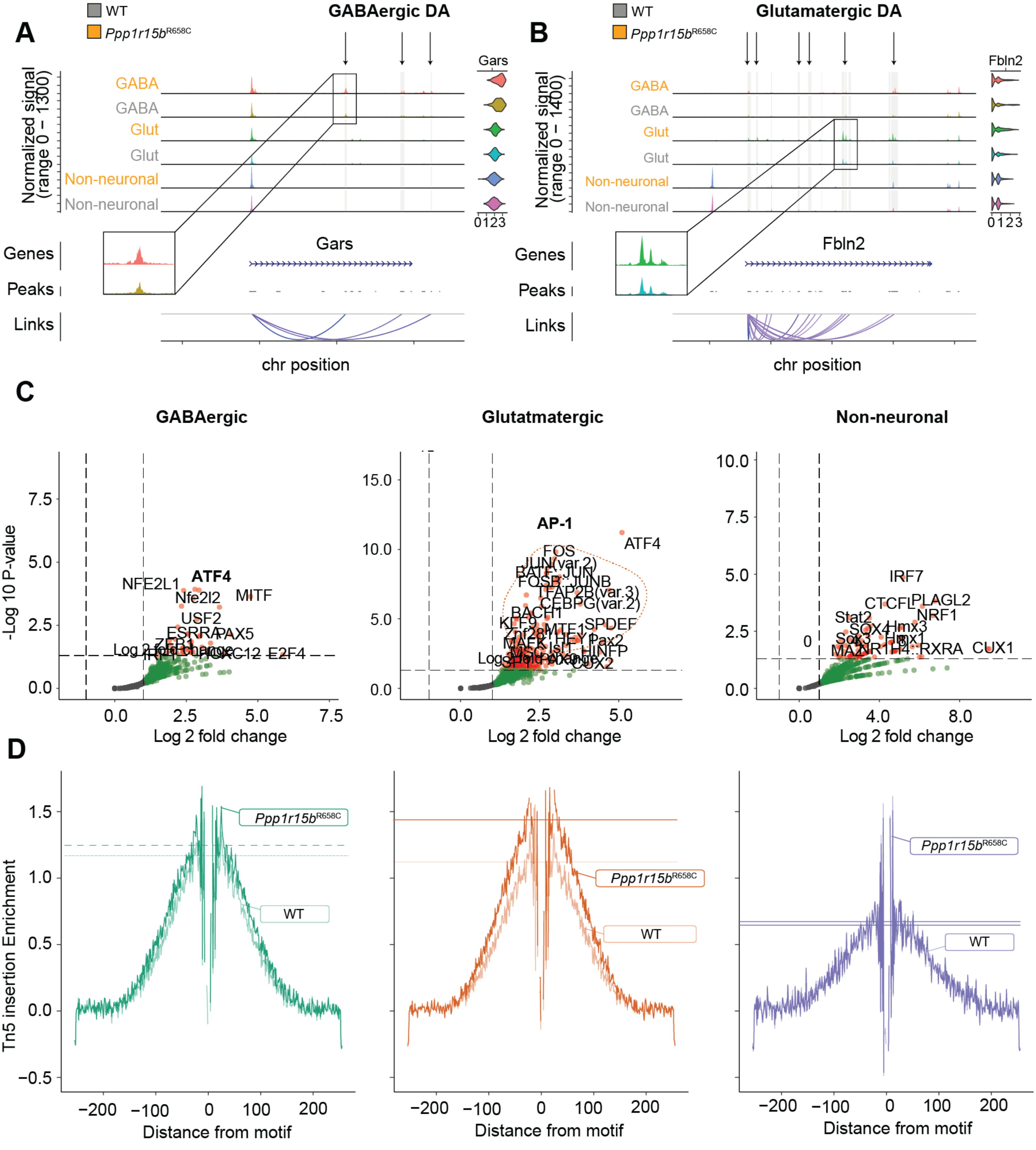


Fig. S7. Torkenczy *et al.*

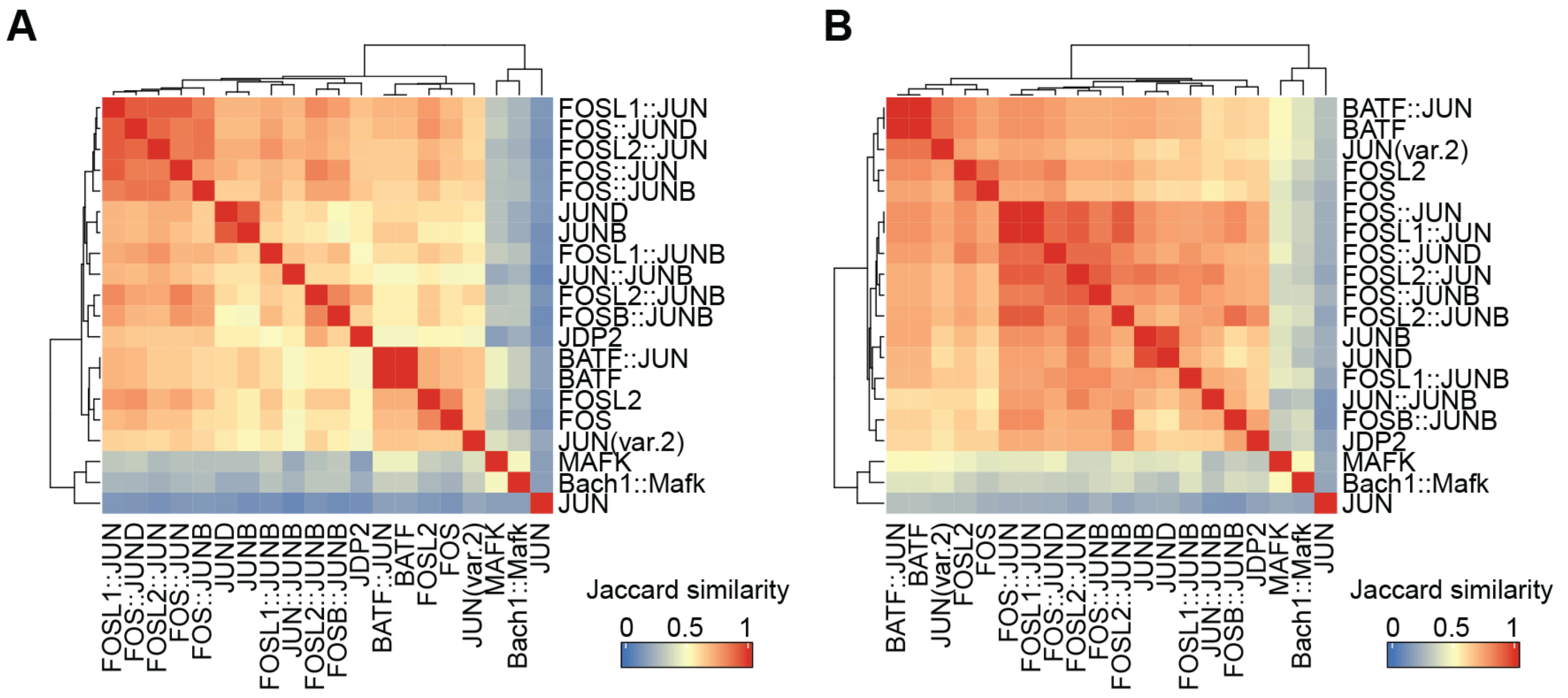


**Fig. S8.** Torkenczy *et al.*

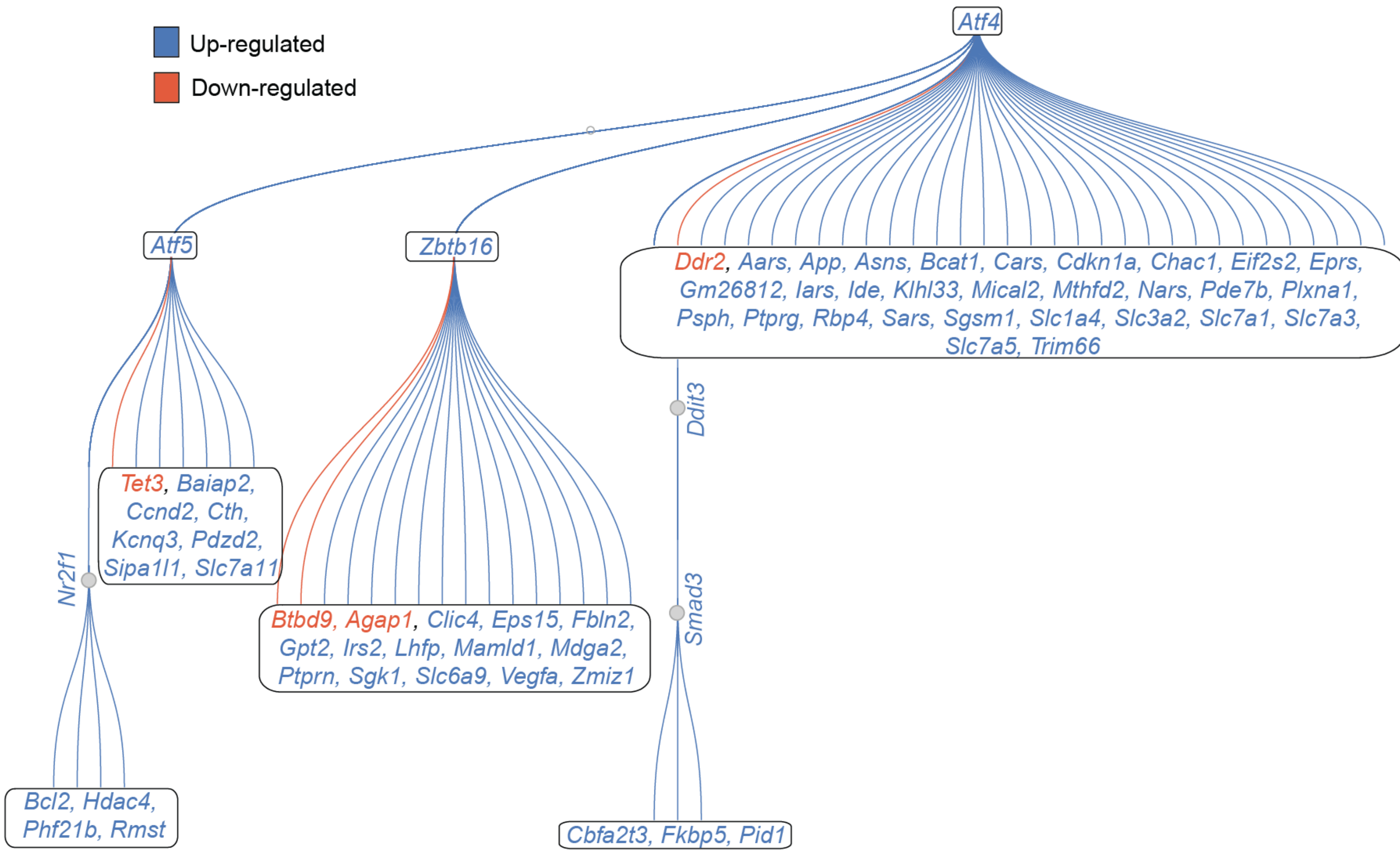


**Fig. S9.** Torkenczy *et al.*

### Supplementary Figure Legends

**Figure S1. Quality control of single-cell sequencing.**

(**A**) Violin plots showing RNA (top panels) and ATAC quality metrics (bottom panels). From left to right: RNA metrics include the number of unique reads per cell, the number of detected genes per cell, and percentage of mitochondrial genes detected per cell. ATAC metrics include the number of peaks detected per cell, the enrichment of Transcription Start Sites (TSS), and the nucleosome banding pattern.

(**B**) Fragment count distribution broken down by fragment length for each sample.

(**C**) UMAP of scRNA-seq data with I3 cell annotations.

(**D**) UMAP of scATAC-seq data with I3 cell annotations.

(**E**) MiloR analysis for differential abundance. Left panel: differential abundance of cell neighborhoods projected into RNA UMAP space. Right panel: cell neighborhoods, split by major cell classes (GABAergic, glutamatergic, and non-neuronal lineages). Dot size indicates the number of cells within a neighborhood, line thickness reflects the number of shared cell types between neighborhoods, and color represents the log fold change in abundance between WT and *Ppp1r15b*$^{R658C}$ cells within each neighborhood.

**Figure S2. Representative differentially expressed genes across cell types and enrichment analysis of molecular function and biological process.**

(**A**) Violin plots showing differentially expressed genes (DEGs) between WT and *Ppp1r15b*$^{R658C}$ cell populations, stratified by cell type. The Kruskal-Wallis test (two-tailed) was used to assess significant increases in ISR genes, with significance indicated by *$P < 0.05$, **$P < 0.01$.

(**B**) Molecular function gene set enrichment analysis results for DEGs. Significant terms were selected with an adjusted *P*-value threshold of < 0.05. Dot color represents the normalized enrichment score (NES), while dot size reflects the $-\log_{10}$ of the adjusted *P*-value.

(**C**) Biological Process enrichment analysis for DEGs. Significant terms were selected with an adjusted *P*-value threshold of < 0.05. Dot color represents the normalized enrichment score, and dot size represents the $-\log_{10}$ of the adjusted *P*-value.

**Figure S3. Comparison of standard single-cell analysis and cluster-free analysis of the ISR upregulated gene set.**

(**A**) Dot plot showing expression of the ISR-specific upregulated gene set across L3 cell types using single-cell normalized expression. The percent of cells expressing each gene is indicated by the dot size.

(**B**) Dot plot showing expression of the ISR-specific upregulated gene set across L3 cell types using a metacell-based aggregation of normalized expression. The percent of metacells expressing each gene is indicated by the dot size.

**Figure S4. Neighborhood analyses of ISR-related changes in the transcriptome.**

(**A**) Iterative analysis of the number of significant neighborhoods required for a gene to be included in Weighted Gene Network Analysis (WGCNA). The number of modules identified in each iteration is shown on the y-axis.

(**B**) Number of genes included in the ISR upregulated gene set (top) and the ISR downregulated gene set (bottom) as a function of the increasing significance cutoff for neighborhood inclusion.

(**C**) U-cell enrichment scores of identified ISR upregulated gene set across cell types. Statistical significance is indicated by (***P* < 0.01, *****P* < 0.0001).

(**D**) U-cell enrichment scores of ISR downregulated gene set across cell types. Statistical significance is indicated by (*****P* < 0.0001).

(**E**) Bar plot showing genes ordered by participation in ISR gene modules across iterations. Genes are color-coded in the ISR upregulated gene set, ISR downregulated gene set, and oligodendrocyte-specific co-regulated gene modules.

**Figure S5. ISR gene set neighborhoods in MiloDE and enrichment across different cell types and brain regions.**

(**A**) ISR upregulated gene set neighborhoods identified using MiloDE, with average fold-change between *Ppp1r15b*$^{R658C}$ and WT cells per neighborhood. Positive fold-change values indicate up-regulation of the module in *Ppp1r15b*$^{R658C}$ cells, while negative values indicate downregulation. Neighborhood size is represented by node size, and overlaps between neighborhoods are shown by edges.

(**B**) ISR upregulated gene set enrichment across I3 cell type annotations.

(**C**) ISR downregulated gene set neighborhoods from MiloDE, with average fold-change between *Ppp1r15b*$^{R658C}$ and WT cells per neighborhood. Positive fold-change values indicate up-regulation in *Ppp1r15b*$^{R658C}$ cells, while negative values indicate down-regulation. Neighborhood size is represented by node size, and overlaps between neighborhoods are shown by connecting edges.

(**D**) ISR downregulated gene set enrichment across I3 cell type annotations.

(**E**) Oligodendrocyte-specific gene set from MiloDE, with average fold-change between *Ppp1r15b*$^{R658C}$ and WT cells per neighborhood. Positive fold-change values indicate upregulation in *Ppp1r15bR658C* cells, while negative values indicate downregulation. Neighborhood size is reflected in node size, and overlaps between neighborhoods are shown by edges.

(**F**) Oligodendrocyte-specific gene set enrichment across I3 cell type annotations.

**Figure S6. Expression patterns of the ISR upregulated module in external datasets.**

(**A**) Dot plot showing the expression of the ISR upregulated gene set across brain cell types in Down Syndrome patients. The average log fold change is represented by dot color, and the -$\log_{10}$ adjusted *P*-value is indicated by dot size.

(**B**) Dot plot showing the expression of the ISR upregulated gene set across brain cell types in Parkinson's Disease patients. The average log fold change is represented by dot color, and the -$\log_{10}$ adjusted *P*-value is indicated by dot size.

(**C**) Dot plot showing the expression of the ISR upregulated gene set across brain cell types in Lewy body dementia patients. The average log fold change is represented by dot color, and the -$\log_{10}$ adjusted *P*-value is indicated by dot size.

(**D**) Dot plot showing the expression of the ISR upregulated gene set across brain cell types in aging mice. The average log fold change is represented by dot color, and the -$\log_{10}$ adjusted *P*-value is indicated by dot size.

**Figure S7. Epigenetic regulation of cell type-specific ISR modulation.**

(**A**) Linked ATAC peaks regulating *Gars* expression (left) and mRNA expression (right) across brain cell types, demonstrating differential accessibility and expression in GABAergic neurons. ±20 kbs from the transcription start site (TSS) is shown. Differential accessibility is indicated by grey shading.

(**B**) Linked ATAC peaks regulating *Fbln2* expression (left) and mRNA expression (right) across brain cell types, demonstrating differential accessibility and expression in glutamatergic neurons. ±20 kb from the TSS is shown. Differential accessibility is indicated by grey lines.

(**C**) Volcano plots showing enrichment of transcription factor motifs in linked regulatory regions for GABAergic (left), glutamatergic (middle), and non-neuronal (right) cell types. A hypergeometric test was performed using GC-content matched background peaks (*P*-value < 0.001 and z-score > 0).

(**D**) Bias-corrected ATAC-seq footprinting signal centered around *FOSL2* motifs, aggregated from peaks linked to gene expression changes across cell types. Dotted horizontal lines represent the flank height for *Ppp1r15bR658C* and WT for comparison.

**Figure S8. Comparison of DNA binding motifs across the AP1 transcription factor family.**

(**A**) Cosine similarity of accessibility of regions regulated by each transcription factor within the AP1 family of transcription factors.

(**B**) Cosine similarity of genes regulated by each transcription factor within the AP1 family of transcription factors.

**Figure S9. Pando analysis supports a role for ATF4 and other transcription factor in the persistent ISR signature.** Hierarchy of regulation centered around ATF4 using Pando gene regulatory networks.
